# TNFR2 blockade promotes anti-tumoral immune response in PDAC by targeting activated Treg and reducing T cell exhaustion

**DOI:** 10.1101/2024.01.22.573571

**Authors:** A. Debesset, C. Pilon, S. Meunier, O. Bonizec, W. Richer, A. Thiolat, C. Houppe, M. Ponzo, J. Magnan, P. Caudana, Jimena Tosello Boari, Sylvain Baulande, N.H. To, B.L. Salomon, E. Piaggio, I. Cascone, J.L. Cohen

## Abstract

**Background:** Pancreatic ductal adenocarcinoma (PDAC) is one of the most aggressive cancers, highly resistant to standard chemotherapy and immunotherapy. Regulatory T cells (Tregs) expressing TNFα receptor 2 (TNFR2) contribute to immunosuppression in PDAC. Treg infiltration correlates with poor survival and tumor progression in PDAC patients. We hypothesized that TNFR2 inhibition using a blocking monoclonal antibody (mAb) could shift the Treg-effector T cell balance in PDAC, thus enhancing anti-tumoral responses.

**Method:** To support this hypothesis, we first described TNFR2 expression in a cohort of 24 PDAC patients from publicly available single-cell analysis data. In orthotopic and immunocompetent mouse models of PDAC, we also described the immune environment of PDAC after immune cell sorting and single-cell analysis. The modifications of the immune environment before and after anti-TNFR2 mAb treatment were evaluated as well as effect on tumor progression.

**Results:** PDAC patients exhibited elevated TNFR2 expression in Treg, myeloid cells and endothelial cells but low level in ductal cells. By flow cytometry and single cell RNAseq analysis, we identified two Treg populations in orthotopic mouse models: resting and activated Tregs. The anti-TNFR2 mAb selectively targeted activated tumor-infiltrating Tregs, reducing T cell exhaustion markers in CD8^+^ T cells. However, anti-TNFR2 treatment alone had limited efficacy in activating CD8^+^ T cells and only slightly reduced the tumor growth. The combination of the anti-TNFR2 mAb with agonistic anti-CD40 mAbs promoted stronger T cell activation, tumor growth inhibition, and improved survival and immunological memory in PDAC-bearing mice.

**Conclusion:** Our data suggest that combining a CD40 agonist with a TNFR2 antagonist represents a promising therapeutic strategy for PDAC patients.

## Introduction

Pancreatic ductal adenocarcinoma (PDAC) is the second leading gastrointestinal cancer in incidence in France. The overall 5-year-survival rate for this malignancy is less than 12% and has not evolved in the last 10 years. In addition, incidence of PDAC is consistently increasing and is expected to become the second leading cause of cancer death in the United States by 2030 [1]. Currently, there is no curative treatment for advanced PDAC. To date, surgical resection offers the only curative option, restricted to solely a small fraction of 20% of patients harboring a localized tumor at the time of diagnosis [2]. For the vast majority of patients diagnosed with advanced or metastatic PDAC, chemotherapy protocols FOLFIRINOX or gemcitabine plus nab-paclitaxel association provide the only available options [3]. PDAC treatment thus remains a true challenge in crucial need for new therapeutic development [4]. In recent years, immunotherapy has emerged as a new tool in cancer treatment. Especially immune checkpoint inhibitors targeting PD-1/PD-L1 axis and the CTLA-4 molecule showed positivive results in melanoma and non small cell lung cancer (NSCLC). Unfortunately, such strategies have shown little to no efficacy in PDAC [5]. The resistance of PDAC is partly due to the intrinsic resistance of cancer cells to therapy, their adaptive plasticity and the tumor microenvironment (TME) [6]. PDAC stroma is composed by a high proportion of cancer-associated fibroblasts (CAFs) and a fibrotic matrix that, together with tumor cells, promote tumor progression, chemoresistance and immunosuppressive signals [7]. TME is characterized by poor infiltration of T lymphocytes [4] [7]. Moreover, the highly fibrotic and hypoxic stroma of PDAC creates a physical barrier to cytotoxic T cells infiltration [8] and promotes immune suppressive cells accumulation such as tumor associated macrophages (TAMs), myeloid-derived-suppressor-cells (MDSCs) and regulatory T cells (Tregs) thus contributing to immune evasion [7,9].

An increased abundance of Tregs has been observed in PDAC tissues and associated with poor prognosis and decreased survival [10–12]. In particular, poor T cell infiltration and Treg enrichment have been described as “immune escape” phenotype and can affect prognostic of PDAC [13,14]. Notably, Treg cell ablation in a mouse model of PDAC resulted in the control of tumor growth dependent on cytotoxic CD8^+^ T cells (CTL) [15]. Tregs highly express the tumor necrosis factor α (TNFα) receptor 2 (TNFR2), and the TNFα/TNFR2 axis was shown to play a crucial role in Treg stability, expansion and function in both mice and humans [16–20]. Moreover, an accumulation of TNFR2^+^ Tregs with superior suppressive capacities have been found across a variety of cancers [21]. Hence, blocking the TNFα/TNFR2 pathway has emerged as a new immunotherapy strategy to target Tregs in cancer [22,23]. In addition, TNFR2 blockade was shown to not only inhibit Treg proliferation but also tumor cells expressing TNFR2 in ovarian cancer and Sezary syndrome [24,25].

Anti-TNFR2 treatment in preclinical experiments has already shown to decrease the frequency of tumor infiltrated Tregs and promote CD8^+^ T cell infiltration and IFNγ expression in subcutaneous mouse models of breast and colon tumors [26,27]. Here, we tested whether blockade of the TNFR2 pathway in PDAC could reverse the balance between regulatory T cells and effector T cells and trigger an effective antitumor response.

## Material and Methods

### Cell culture

Murine pancreatic cancer cells mPDAC, were isolated and validated, as previously described [28] from tumor-bearing *p48cre, Kras^LSL_G12D^, p53^R172H/+^, Ink4a/Arf^flox/+^* FVB/n mice (kindly provided and validated by D. Saur, TranslaTUM, Munich, Germany), and were cultured in DMEM (41965-039, Gibco) 10% FBS. Murine pancreatic cancer cell line Panc-02 (kindly provided by R. Ronca (University of Brescia) in 2016) were cultured in RPMI 10% FBS and pyruvate 0,1%.

### Cell viability assay

A total of 5×10^3^ mPDAC cells/well were plated in 96-multiwell plates. The day after, the cells were treated with 12.5, 25 or 50 µg/mL of anti-TNFR2 mAb or IgG control. After 72 h, cell viability was analyzed by Alamar Blue Assay, as previously described [28].

### Tumor mouse model

Female FVB/n or C57BL/6j mice of 10 to 14 weeks of age were obtained from Janvier Labs (France). All *in vivo* experiments were carried out with the approval of the appropriate ethical committee (authorization number #24225-202001310859869 and #29529- 2020110222005935) and under conditions established by the European Union. All experiments were performed in the same animal facility (IMRB).

#### For tumor orthotopic mouse models

10 to 14 weeks of age FVB/n or C57BL/6j mice were injected orthotopically in the pancreas with mPDAC cells (10^3^ cells/mouse in 50µL) or Panc02 (10^5^ cells/mouse in 50µL) respectively, as previously described [28].

#### For tumor ectopic model

10 to 14 weeks of age FVB/n mice were injected subcutaneously with mPDAC cells (105cells/mouse in 100 µL) on the right flank.

### Antibody treatments

Polyclonal Armenian hamster IgG (ref: BE0091), anti-mouse-TNFR2 (clone TR75-54.7, ref: BE0247), anti-mouse CD40 agonist (clone FGK4.5/FGK45, ref: BE0016-2), anti-mouse-CD8 alpha (clone 53-6.7, ref: BP0004) monoclonal antibodies (mAbs) were purchased from BioXCell (Euromedex, France). After orthotopic engraftment, FVB/n and C57Bl/6j mice received 3 intraperitoneal injections of anti-TNFR2 mAb (500 µg in100 µL) on days 11, 13, and 15 or 5 injections at days 8, 11, 13, 15 and 18 respectively. Before treatments, animals were randomised in cages and the same investigator did the measures. For combination therapy, mice received 3 intraperitoneal injections of CD40 agonist mAb (100 µg in 100 µL) on days 11, 13, and 15. Control groups received either IgG control (500μg), PBS or were untreated (as indicated on figures’ legends). Potential cofounders as order administration of the therapeutic molecules were not evaluated. After sub-cutaneous engraftment (Supplementary Figure 1), FVB/n mice received 3 intraperitoneal injections of anti-TNFR2 mAb or IgG control (500 µg in100 µL) on days 8, 11, and 13. Tumor growth was monitored after reaching a volume of approximately 150-200 mm^3^. Mice were euthanized at day 18 or 21 after orthotopic engraftment (as indicated on figures’ legends) or day 20 after subcutaneous engraftment. At this time, total tumor burden was quantified as previously described [28], tumors and draining lymph nodes were collected for flow cytometry analysis.

*For survival experiment*, FVB/n (mPDAC) and C57Bl/6j (Panc02) mice received anti-TNFR2 and/or CD40 agonist on days 11, 13, 15, 20, 27 and 34 after orthotopic engraftment. Control groups received PBS. Mice were euthanized when the clinical score reached a limit established by a grid of symptoms (ventral swelling, slimming, anemia and cachexia). After 64 days, tumor presence in C57Bl/6j (Panc02) was assessed by echography. After 81 days, mice still alive and naïves mice were injected subcutaneously with mPDAC cells (10^5^cells/mouse in 100 µL) on the right flank and tumor growth was monitored.

### Flow cytometry analysis

Lymph nodes were mechanistically dissociated, counted and passed through 70 µm cell strainers (Falcon 352350). To harvest infiltrated immune cells, tumors were minced with scissors and digested in RPMI-1640 2%-FBS, Collagenase IV [367,5 U/mL] (Worthington Biochemical, ref: WOLS04186) and DNase I [0,1 mg/mL] (Sigma, ref D5025) at 37 °C for 60 min under rotation. Digested tumors were passed through a 70 mm cell strainer, and centrifuged at 400×g for 5 min. Then, cell suspension was centrifuged in a gradient of Ficoll (GE Healthcare, ref: 17144003) for 40 min at 400g. All samples were stained as previously described [29] with the antibodies listed in Supplemental Material. Non-specific binding was blocked using anti-CD16/CD32 (Miltenyi, ref: 130-07-594). For cytokines staining, cells were stimulated in RPMI 10% FBS, PMA [1μg/ml] and ionomycin [0,5μg/ml] with Golgi Plug [1:1000] and Golgi Stop [1:1000] solutions (Cytofix/Cytoperm TM Plus, BD Bioscience, ref: 555028) for 4-5h at 37°C to block Golgi’s exocytosis. For intracellular staining, cells were fixed and permeabilized with fixation/permeabilization (Invitrogen, ref: 00-552300) according to the manufacturer’s instructions. Data were acquired with a BD Bioscience Canto II or a BD LSR-Fortessa flow cytometer, compensated and exported into FlowJo software (TreeStar Inc.) for analysis. After analysis, the data points <100 events were excluded. The complete list of Abs used can be found in supplementary material.

### Analysis of Single-cell RNA sequencing published data of PDAC patients

The expression of TNFR2 in human PDAC patients was analyzed by using public single cell RNAseq data [30]. Data were analyzed from raw counts matrix using standard Seurat workflow [31]. Briefly, low quality cells (<200 genes/cell, <3 cells/gene and >10% mitochondrial genes) were excluded. “NormalizeData” function with default parameters was applied to normalize the expression level of genes in each single cell. Then, 3,000 highly variable genes were identified using the “FindVariableFeatures” function with ‘vst’ method. All samples were processed independently and the data was then integrated using reciprocal PCA. The “ScaleData” function was used to scale and center gene expression matrices after regressing out heterogeneity associated with mitochondrial contamination. To perform clustering, the dimensionality of the data was determined by calculating relevant principal components using the ElbowPlot function. Relevant principal components were selected to construct the shared nearest neighbor (SNN) graph with “FindNeighbors” function, and clusters were determined using the Louvain algorithm. The uniform manifold approximation and projection (UMAP) was finally applied based on the above described SNN graph to visualize the single cell transcriptional profile in 2D space. Annotation of the clusters was performed using marker genes and published gene signatures. Imputation of missing values in the count matrix was performed using Adaptively thresholded Low-Rank Approximation [32].

### Single-cell RNA sequencing of mouse models

Murine pancreatic cancer cells mPDAC were injected of syngenic immunocompetent FVB/n mice as previously described [28]. Mice were treated with anti-TNFR2 mAb or untreated. After 18 days, mice were euthanized, and tumors were processed with Collagenase IV and DNase I as previously described (flow cytometry). Cells were stained with DAPI, CD45, CD4, CD8, CD11c, Ly6G, Ly6C, CD19, NK1.1, F4/80 and T cells were sorted on a FACSAria III (BD) by gating on live cells, CD45^+^, lineage (CD11c, Ly6G, Ly6C, CD19, NK1.1, F4/80) negative cells, CD4^+^ and CD8^+^ cells. CD4 and CD8 isolated T cells were mixed. Sorted T cells samples were loaded on a 10x Chromium Controller (10X Genomics) according to manufacturer’s protocol. Single-cell RNA-Seq libraries were prepared using Chromium Single Cell 5′ v3 Reagent Kit (10x Genomics) according to manufacturer’s protocol. Briefly, the initial step consisted in performing an emulsion where individual cells were isolated into droplets together with gel beads coated with unique primers bearing 10x cell barcodes, unique molecular identifiers (UMI), and poly (dT) sequences. Reverse transcription reactions were engaged to generate barcoded full-length cDNA followed by the disruption of emulsions. cDNA was then purified using DynaBeads MyOne Silane Beads (Thermo Fisher Scientific). PCR amplification of 10x Barcoded full-length cDNA was performed following manufacturer’s instructions. Finally, libraries were constructed following these steps: (1) fragmentation, end repair and A-tailing; (2) size selection with SPRI select beads; (3) adaptor ligation; (4) post-ligation cleanup with SPRI select beads; (5) sample index PCR and final cleanup with SPRI select beads. Library quantification and quality assessment were achieved by Qubit fluorometric assay (Invitrogen) using dsDNA HS (High Sensitivity) Assay Kit and Bioanalyzer Agilent 2100 System using a High Sensitivity DNA chip. Indexed libraries were tested for quality, equimolarly pooled and sequenced on an Illumina HiSeq2500 using paired-end 26 × 98bp as sequencing mode. By using a full Rapid flow cell, coverage was around 100M reads per sample corresponding to 100,000 reads per cell.

After sequencing, single-cell expression was analyzed using the Cell Ranger Single Cell Software Suite (v3.1.0) to perform quality control, sample de-multiplexing, barcode processing, and single-cell 5′ gene counting. Sequencing reads were aligned on 10xGenomics mm10-3.0.0 mouse genome reference using the Cell Ranger suite with default parameters. This version of Cell Ranger including EmptyDrops method, cells with low RNA content has been rescue. Downstream analyses were performed using Seurat (v4.3.0) with R version 4.3.0. Some filters were applied for each sample imported into seurat pipeline: cells with fewer than 200 genes were removed to remove debris, deaths cells and other cells with few genes. Uninformative cells and possible doublets cells were removed based on the percentage of mitochondrial genes (cells with percentage mitochondrial genes superior to 0,05 are filtered) and total counts of genes by cell (cells with a value of genes/cells inferior to 350 are filtered). Each sample were normalized separately using global-scaling normalization (using NormalizeData function with ‘‘LogNormalize’’ as normalization method) and the 2,000 most highly variable genes were identified for each sample (using FindVariableFeatures with ‘‘vst’’ as method, with low- and high-cutoffs for feature dispersions fixed at 0.5 and Inf and with low- and high-cutoffs for feature means fixed at –Inf and Inf; in addition, Tra[vdj] and Trb[vdjc] genes are filtered from high variable variable genes). All the samples were merged applying the integration process of Seurat based on the 2000 most variable features (using SelectIntegrationFeatures, FindIntegrationAnchors and IntegrateData with default settings). To reduce technical noise, Principal Component analysis (PCA) were performed to work on the most contributing principal components (PC). Graph-based clusterization was done at different resolution (using FindNeighbors on the thirty first PCs and FindClusters for the resolution between 0 and 1 for each decimal) and visualized using Clustree version 0.5.0 (Zappia, Oshloack, 2018). UMAP reduction was done (using RunUMAP on the thirty first PCs) to visualize the cells in UMAP projection.

Clustering with resolution 0.2 was satisfying for the identification of contaminant cell based on absence of expression of T cell markers (Cd3e, Cd3d, Cd4, Cd8) and expression of other immune cell population markers (Cd14). After elimination of contaminant cells from the data of the different sample, the different steps described above (from normalization to UMAP reduction) were performed again.

Cluster identification: 12 clusters were initially defined at resolution 0.7 for all CD3^+^ cells and are visualized using UMAP. Cluster were assigned using “find all markers” analysis and known T cells marker. The expression of a selection of genes was then analyzed using Feature plots and Violin plots. For CD8 cells analysis the “add module score” function was used to compare an exhausted score using Tigit, Havcr2, Ctla4, Lag3 and Tox genes.

### ScTCR-seq analysis

After sequencing, single-cell TCR data were analyzed using the Cell Ranger Single Cell Software Suite (v3.1.0) to perform quality control, sample de-multiplexing, barcode processing, and single-cell VDJ CDR3 counting. Sequencing reads were aligned on 10xGenomics vdj_GRCm38_alts_ensembl-3.1.0.gz-3.1.0 genome reference using the Cell Ranger suite with default parameters.

Filtered files produced by Cellranger were imported in R to study clonality. To improve the definition of clonotypes and considering that TRA and TRB chains are defined by combining CDR3 and V subunits, we considered for our analysis only those cells presenting only 1 TRA chain and 1 TRB chain to define clones. To estimate clonal diversity, we used Gini-TCR Skewing Index [33].

### Statistical analysis

Statistical analyses were performed by using GraphPad Prism software. Unless indicated otherwise, bars represent mean ± Standard Error of Mean (S.E.M.). For flow cytometry, histology analyses, and tumor volume comparison a parametric (Student *t*-test, ANOVA) or a non-parametric (Kruskal-Wallis, Mann-Whitney) test were applied and outliers were excluded. In the experiment comparing anti-TNFR2 toward control mice, Student *t*-test one-tailed was used to test if the anti-TNFR2 was lower than control groups. For Kaplan-Meier survival curves, groups were compared using the log-rank test. Chi-quare test was used to compare the proportion of mice with a tumor. Statistical signifance is indicated as ns: non-significant *p<0,05, **p<0,01, ***p<0,001,****p<0,0001.

## Results

### TNFR2 emerges as a promising novel target in PDAC treatment

Targeting TNFR2 for anti-tumor purposes through the use of anti-TNFR2 mAb is based on the possibility to block local immunosuppression mediated by Treg constitutively expressing TNFR2 and by the putative expression of this marker directly by certain tumor cells [24]. We first took the opportunity of available public single cell RNAseq data [30] to evaluate the expression of the TNFR2 gene (*Tnfrs1B*) in PDAC patients. Clusters of the main populations present in PDAC tumors were assigned using module score of lists of differentially expressed genes published in Peng et al. [30] and visualized by UMAP (**Figure 1A**). The level of expression of *Tnfrsf1b* among the different main cell populations identified in PDAC was then analyzed. As expected, the main cell populations expressing *Tnfrsf1b* in the tumors are regulatory T cells, myeloid and endothelial cells (**Figure 1B**). Importantly, after separating healthy (n=11) and tumour (n=24) single cell RNAseq data, the immune infiltration radically differed (**Figure 1B**). Whereas conventional T cells and Tregs were not detected in healthy pancreas, they were present in tumour environment and they highly expressed *Tnfrsf1b* (**Figure 1B and C**). Among epithelial-derived cells, the ductal 2 cell clusters were present only in tumors and were characterized by markers for classical and basal-like PDAC cells [34,35]. Ductal 2 cell clusters expressed *Tnfrsf1b* at low level (**Figure 1A and B)**. Stromal cells and ductal 1 cells expressed very low level of *Tnfrsf1b* both healthy and tumour tissues (**Figure 1A and B**). These results support the relevance of TNFR2 as a potential target in immune cell infiltrate of human PDAC.

**Figure 1.**
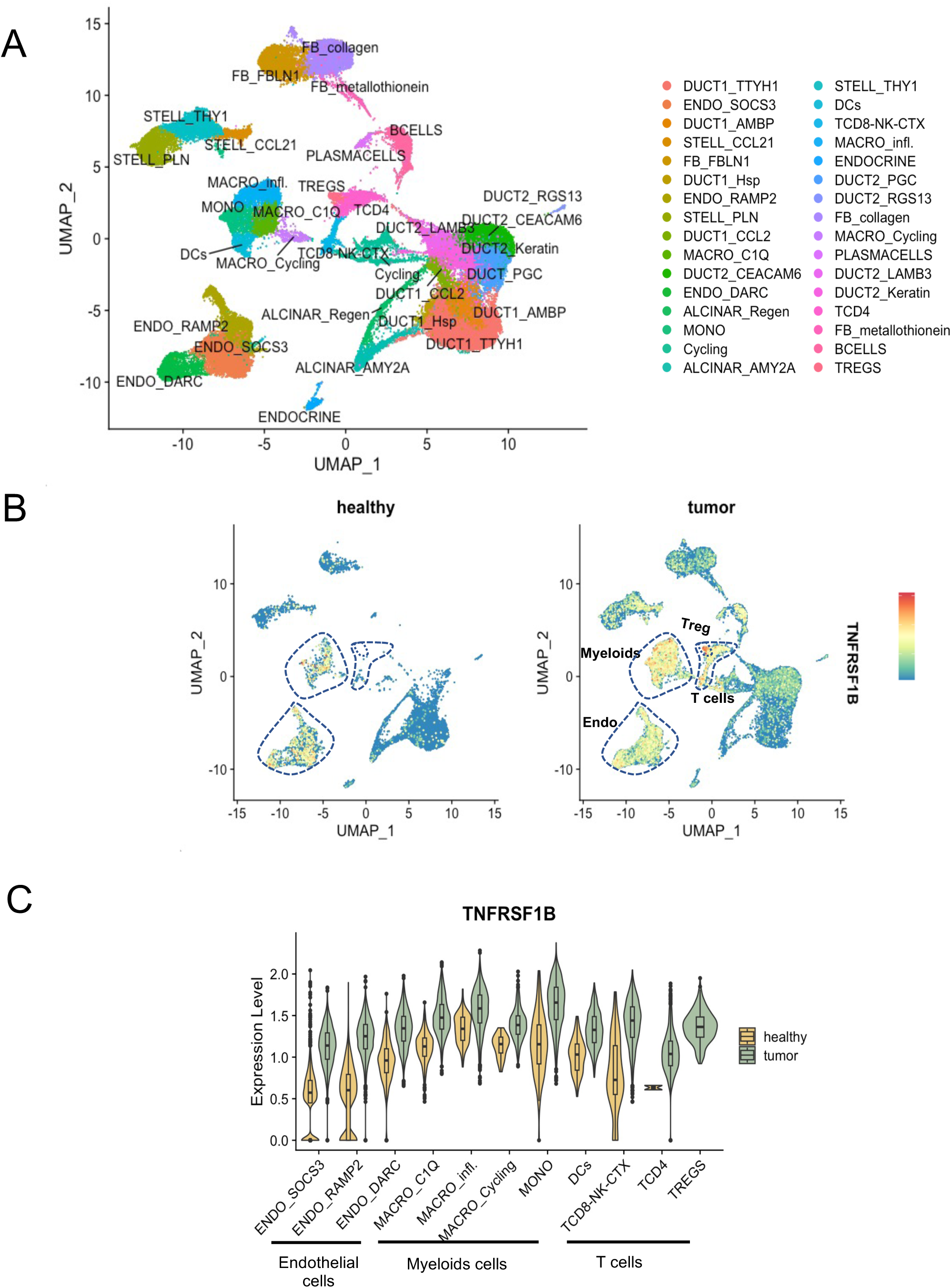
TNFR2 expression in human PDAC. (**A**) UMAP plot of main populations present in PDAC tumors from Peng et al [30]. (**B**) Expression of TNFRSF1B split between tumor and healthy samples. (**C**) Violin plot displaying the expression of TNFRSF1B across the cell identified in PDAC. Adaptive thresholded low rank approximation-imputed data are depicted.

### TNFR2 blockade decreases tumor-infiltrating Treg proportions in PDAC tumors

We sought to study the expression of TNFR2 by tumor infiltrated immune cells in orthotopic and immunocompetent mouse models of PDAC obtained by using mPDAC and Panc02 cell lines previously described [28]. We initially observed that mPDAC tumors have an immune-escape TME infiltrated by CD45^+^ immune cells, higly enriched in CD4^+^Foxp3^+^ Tregs and characterized by low CD8^+^ T cell infiltration [29]. Here, tumor infiltrated lymphocytes (TILs) were isolated from mPDAC tumors and analyzed by flow cytometry. 14% (± 3) of CD45^+^ infiltrated immune cells were CD3^+^ T cells and 69% (± 3) were CD11b^+^ myeloid cells (**Figure 2A**). TNFR2 was detected in CD3^+^ T cells, CD11b^+^ myeloid cells (**Figure 2A**) and in tumororal cells (**Figure 2B**). Among T cells, the intensity of TNFR2 expression was significantly higher in tumor infiltrated CD4^+^Foxp3^+^ Tregs than in CD4 conventional (CD4conv) CD4^+^Foxp3^-^ T cells or CD8^+^ T cells (**Figure 2C**). The vast majority of Tregs infiltrating the mPDAC tumor express TNFR2 but lower proportion of CD8^+^ (38%) and CD4^+^Foxp3^-^ T cells (21%) expressed this receptor (**Figure 2C**). Comparable results of higher TNFR2 expression in tumor infiltrated CD4^+^Foxp3^+^ Tregs compared to CD4^+^Foxp3^-^ T cells were observed in a second model of orthotopic PDAC tumors obtained by injecting murine Panc02 cells (**Supplementary Figure S1A-C**).

**Figure 2.**
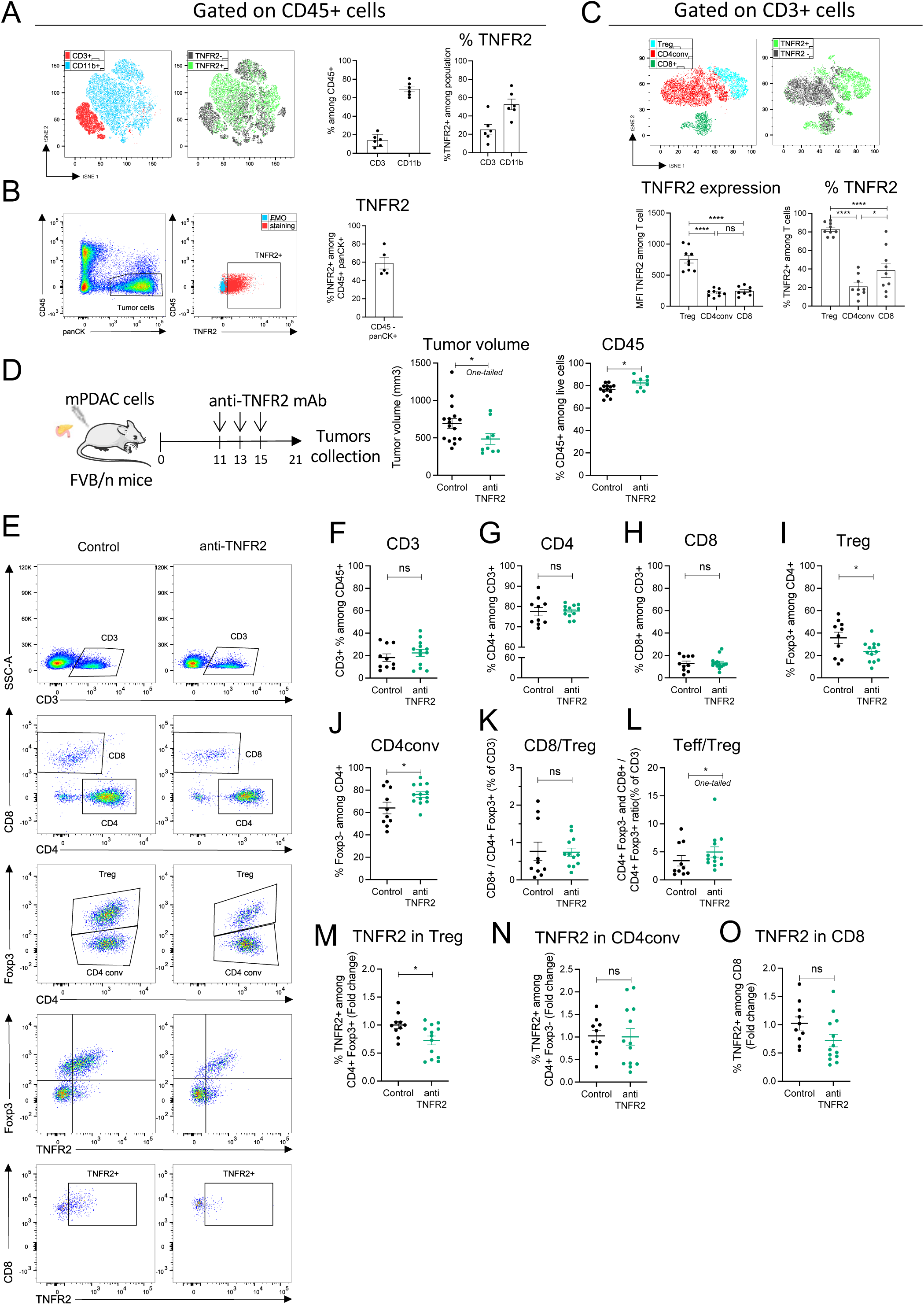
TNFR2 blockade decreases tumor volume and tumor-infiltrating Treg proportions in mouse models of PDAC. FvB/n immunocompetent mice were grafted with mPDAC cells into the pancreas. After 21 days, tumors were harvested for flow cytometry analysis. **(A)** Cell clustering using a t- distributed stochastic neighbor embedding (t-SNE) algorithm performed on CD45^+^ FVS- (Fixable Viability Stain) gate. CD3^+^ and CD11b^+^ cell clusters localization and TNFR2^+^ cell localization among the above-mentioned populations are indicated. Histograms show the proportion of intratumoral CD3^+^ and CD11b^+^ cells among CD45^+^ FVS- cells and of TNFR2^+^ cells among CD3^+^ or CD11b^+^ cells. **(B)** Gating strategy of tumor cells (CD45^-^ panCK^+^) and TNFR2^+^ cells among tumor cells and proportion of TNFR2^+^ cells among CD45^-^panCK^+^ cells. (**C)** Cell clustering using a t-SNE algorithm performed on CD3^+^ gate. Treg (CD4^+^Foxp3^+^), CD4conv (CD4^+^Foxp3^-^) and CD8^+^ cell clusters, and TNFR2^+^ cell cluster localization are indicated. Histograms show TNFR2 expression (MFI TNFR2) and proportion in intratumoral Treg (CD4^+^Foxp3^+^), CD4conv (CD4^+^Foxp3^-^) and CD8^+^ cells. **(D)** Schemas of the experiment: mice were treated with anti-TNFR2 mAb, IgG control or PBS at day 11, 13 and 15 or were untreated. Tumor volume at day 21 was measured and infiltrated CD45^+^ cells among FVS- cells analyzed without previous separation on density gradient (n=16). **(E)** Gating strategy of infiltrated T lymphocytes analyzed by flow cytometry. Representative scatter plots of intratumoral (**F-H**) CD3^+^, CD4^+^ and CD8^+^; (**I**, **J**) Treg (CD4^+^Foxp3^+^) and CD4conv (CD4^+^Foxp3^-^); **(K-L)** CD8^+^/Treg(CD4^+^Foxp3^+^) ratio, and Teff (CD4^+^Foxp3^-^ and CD8^+^)/Treg(CD4^+^Foxp3^+^) ratio. **(M-O)** TNFR2 proportion among Treg (CD4^+^Foxp3^+^), CD4conv (CD4+ Foxp3-) and CD8+ cells is represented as fold change (calculated by reporting each point to the mean of the control group). For (**F-O**) (n=20 including 2 experiments, representative of 3 experiments). Data are plotted as the mean ± SEM. Statistical significance between population in control was determined using ANOVA test. Statistical significance from controls was determined using a t-test [F-G-I-J-N-O-P] or Mann-Whitney test (two-tailed [H-K-R] or one-tailed [L-M] (depending of the data following a normal distribution). Ns: non-significant, p>0,05, *p<0,05, **p< 0,01, ***p<0,001.

Since disruption of the TNFα/TNFR2 pathway was shown to decrease tumor growth and metastasis progression in different experimental models of cancer [21], we sought to test the effect of TNFR2 blockade on PDAC progression and on the regulation of the immune microenvironment in the mPDAC and Panc02 orthotopic or ectopic tumors. We previously observed a Treg enrichment between day 7 and day 17 in mPDAC tumor bearing mice [29]. To target CD4^+^Foxp3^+^ Tregs expressing TNFR2, mice were treated with 3 injections of a blocking anti-TNFR2 monoclonal antibody (mAb) at day 11, 13 and 15 for mPDAC tumors (**Figure 2D**) and at day 8, 11, 13, 15 and 18 for Panc02 tumors (**Supplementary Figure S1D**). 21 days after mPDAC or Panc02 cell injection, mice were euthanized and their tumor volumes measured. The mPDAC and Panc02 tumor volumes statistically decreased in mice treated with the anti-TNFR2 (**Figure 2D and Supplementary Figure S1E**). TNFR2 blockade did not have any effect on the progression of mPDAC ectotopic tumors (**Supplementary Figure S1L and M**). We analyzed the TILs after anti-TNFR2 mAb administration by flow cytometry. The frequency of CD45^+^ among total number of cells isolated from tumor digestion increased in anti-TNFR2 treated mice (**Figure 2D**). The proportion of CD3^+^, CD4^+^ and CD8^+^ cells among CD3^+^ cells was not modified (**Figure 2E-H and Supplementary Figure S1F**). However, the frequency of CD4^+^Foxp3^+^ Tregs cells significantly decreased by 1.5 fold both in mPDAC and Panc02 tumor-bearing mice (**Figure 2I and Supplementary Figure S1G**) whereas the CD4^+^Foxp3^-^ (CD4conv) increased (**Figure 2J**). While the CD8^+^/Treg ratio did not change (**Figure 2K and Supplementary Figure S1I**), the ratio between all effector T cells (CD4^+^Foxp3^-^ and CD8^+^/CD4^+^Foxp3^+^) and Treg (Teff/Treg) increased (**Figure 2L and Supplementary Figure S1J)** in anti-TNFR2 treated mice compared to untreated mice. Also, the frequency of TNFR2 among CD4^+^Foxp3^+^ cells but not of CD4^+^Foxp3^-^ or CD8^+^ T cells was lower in anti-TNFR2 treated mice compared to untreated mice (**Figure 2M-O**). In tumor draining lymph nodes (TDLN), no major modification were observed (**Supplementary Figure S2**). We did not observe any impact of the treatment on the immune infiltrate of mPDAC when tumors were ectotopically injected (**Supplementary Figure S1K-N**), in accordance with the absence of clinical impact of anti-TNFR2 treatment in the last setting.

Since, TNFR2 was also expressed by CD11b^+^ infiltrating the tumors (**Figure 2A**), TNFR2 expression and blocking effect were analyzed in different types of myeloid cells (**Supplementary Figure S3A**). TNFR2 was highly expressed by the tumor associated macrophages (TAMs) and the polarized macrophages M1 and M2 CD206^+^-cells (**Supplementary Figure S3B-D**). Also, it was shown that TNFα drives the accumulation of peripheral MDSCs via TNFR2 signaling, regulating MDSC survival and helps tumor cells evade from the immune system [36]. In mPDAC mouse models, infiltrated CD11b^+^ suppressive cells constituted more than 50% of the CD45^+^ cell populations as in the human pathology [29]. PMN-MDSCs directly inhibit T cell proliferation, express PDL-1 and immunosuppressive cytokines and mobilizes Treg [37]. Indeed, PMN-MDSCs are the dominant source of TNFα leading to stromal inflammation and immune tolerance to promote therapeutic resistance in PDAC [38]. This was observed by using the etanercept, a soluble form of TNFR2, to block TNFα. However, etanercept inhibits both TNFR1 and TFR2 signaling by targeting TNFα. In mice treated with anti-TNFR2, the proportion of the CD11b^+^ myeloid cells was not modified (**Supplementary Figure S3E**) whereas the frequency of polymorphonuclear myeloid-derived suppressor cells (PMN-MDSCs) but not the monocytic myeloid-derived suppressor cells (M-MDSCs) significantly decreased (**Supplementary Figure S3F and G**). No impact on the % of TAMs, M1, M2 CD206^+^ or M2 CD206- (**Supplementary Figure S3H-J**) or dendritic cells (**Supplementary Figure S3K-M**) or CD80 and CD86 expression by DC (**Supplementary Figure S3N-P**) was observed. These results indicate that despite a strong expression of TNFR2 on tumor infiltrated myeloid cells, anti-TNFR2 mAb treatment only impact PMN-MDSCs in mPDAC tumors.

Altogether, these results suggest that anti-TNFR2 mAb treatment mainly targeted Tregs and PMN-MDSCs of the pancreatic TME thus promoting higher infiltration of immune cells leading to a more favorable balance beween effector T and Treg cells in the TME.

### TNFR2 blockade decreases Treg activation and effector T cell exhaustion

To go further inside of the mechanisms of regulation due to anti-TNFR2 mAb treatment, we next analyzed mPDAC TILs by single cell RNAseq analysis. For this, mice were euthanized at day 18 to be closer to the end of the anti-TNFR2 mAb treatment, and thus to increase the possibility to detect any effect. We isolated CD45^+^ cells from tumor-control mice and anti-TNFR2-treated mice at day 18. CD3^+^ cells were sorted and analyzed using single cell RNAseq. After data normalization and using known markers of T cells, cells are clustered in two dimensions using the UMAP dimensionality reduction technique (**Supplementary Figure S4**). Two clusters of Treg (activated, (a)Treg and resting (r)Treg) were observed and cell clusters of CD4^+^ effector/memory T cells (*Cd44*, *Cd40L*, *Cxcr6, Icos*), CD8 effector/memory (*Cd8a, Pdcd1*, *Gzmb FasL*), naives CD8^+^ and CD4^+^ (*S1pr1*, *TCF7*, *Lef1*, *Sell*) and cycling cells (*Mki67)* were identified (**Supplementary Figure S4**). Focusing on the Treg clusters, rTreg and aTreg clusters were separated by differently expressed genes such as *Il2ra, Klrg1, Lgals1, Gzmb, Ccr8,* (**Figure 3A**). The activation markers *Pdcd1*, *Ctla4*, *Icos*, *Itgb7* and *Tnfrsf1b* were more expressed in aTreg than in rTreg (**Figure 3B**). Anti-TNFR2 treatment induced deregulation of a greater number of genes in the aTreg (924 genes) cluster than in the rTreg one (423 genes) (**Figure 3C**). In particular, it induced a decrease of several genes of activation and of genes of the NF-Kβ signalling pathway (**Figure 3D**). Importantly, we observed that anti-TNFR2 mAb treatment reduced the expression level of *Tnfrsf1b* and *Ctla4* (**Figure 3D**). We validated these anti-TNFR2 effects by flow cytometry. The frequency of CD4^+^FoxP3^+^ Tregs expressing TNFR2, CTLA4 or both significantly decreased under anti-TNFR2 mAb treament (**Figure 3E-G**). TCR sequencing revaled that among infiltrated T lymphocytes, cell populations displaying high numbers of expanded clones were identified in aTregs, CD4-Eff/mem, cycling and CD8/CTL clusters (**Figure 4A and B**). We next calculated the Gini index which captures the inequality in clonotype size across the population (**Figure 4C**). First, its global value was limited (less that 0.3) suggesting a low clonal expansion in the PDAC tumors. As expected, activated aTreg, CD4-eff/mem and CD8- CTL clusters had the higher Gini index, reflecting the more highly expanded clones. Interestingly, anti-TNFR2 mAb treatment induced Gini index reduction for the 3 first mentioned clusters, with a less pronounced effect for the CD8-CTL cluster (**Figure 4 C**). Importantly, anti-TNFR2 treatment specifically targets aTregs, as evidenced by the strong reduction of the percentage of cells with expanded clones, likely revealing a less immunosuppressive environment in the tumor of treated mice (**Figure 4D**).

**Figure 3.**
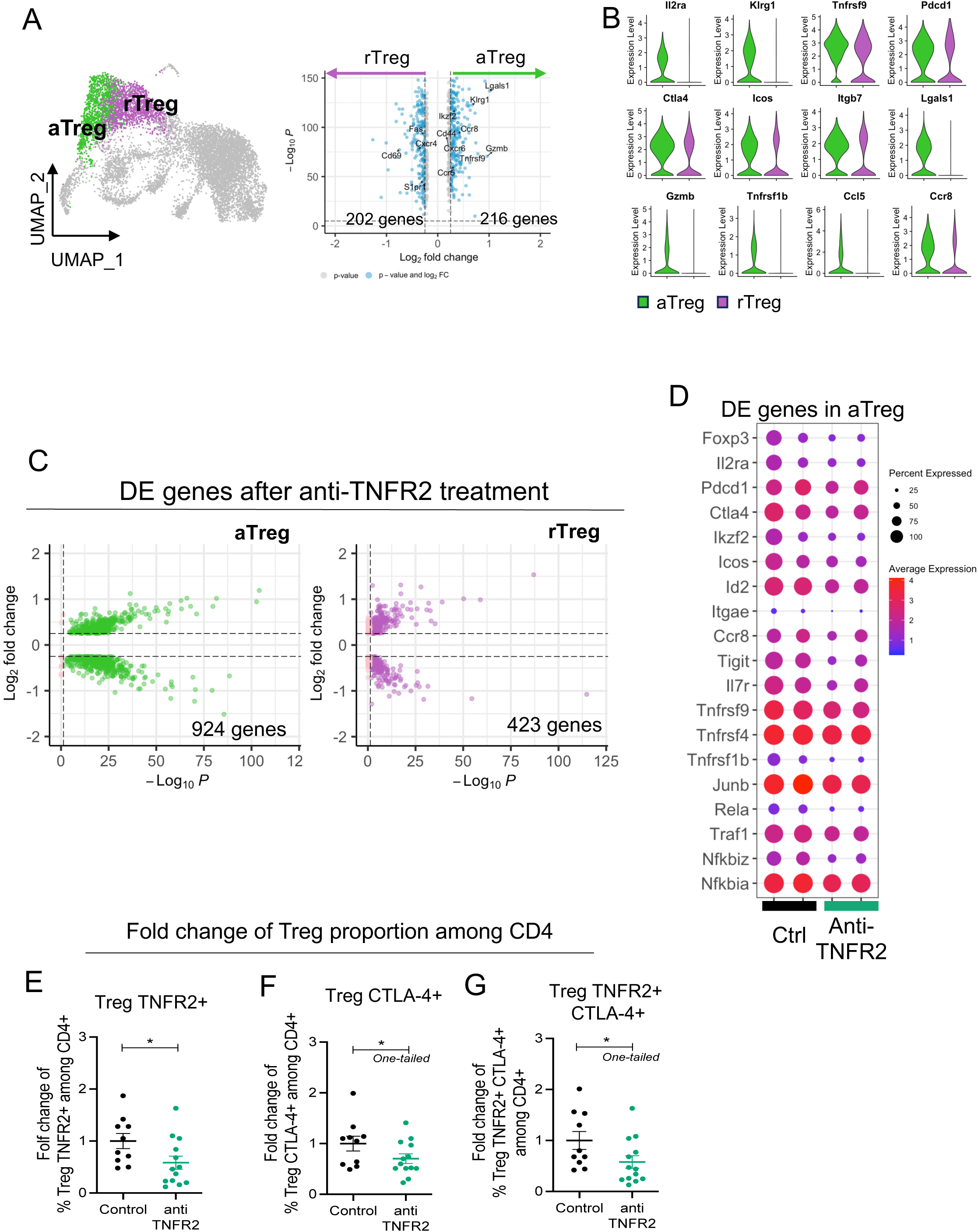
TNFR2 blockade selectively targets activated Tregs over resting Tregs in a PDAC mouse model. FvB/n immunocompetent mice were grafted with mPDAC cells into the pancreas. Mice were treated with anti-TNFR2 mAb and tumors were harvest and intratumoral CD4^+^ and CD8^+^ were isolated for single–cell RNA sequencing. **(A)** UMAP plot with showing the two identified Treg clusters and violin plot comparing expression of different genes between aTreg (green) and rTreg (purple). **(B)** Volcano plot of differential gene expression analysis between aTreg and rTreg. **(C)** Volcano Plot of differential gene expression analysis between control and anti-TNFR2 treated mice for aTreg (Left) and rTreg (right) clusters. **(D)** Dot plot highlighting anti-TNFR2 treatment effect on gene expression level on aTreg. (E-G) Scatter plots of fold change of (E) Foxp3^+^ TNFR2^+^ (Treg TNFR2^+^), (F) Foxp3^+^ CTLA-4^+^ (Treg CTLA4^+^) and (G) Foxp3^+^ TNFR2^+^ CTLA-4^+^ (Treg TNFR2^+^ CTLA-4^+^) proportions among CD4. Data are plotted as the mean±SEM. Statistical significance[from controls was determined using a Mann-Whitney test two-tailed or one-tailed [F-G], *p<0,05.

**Figure 4.**
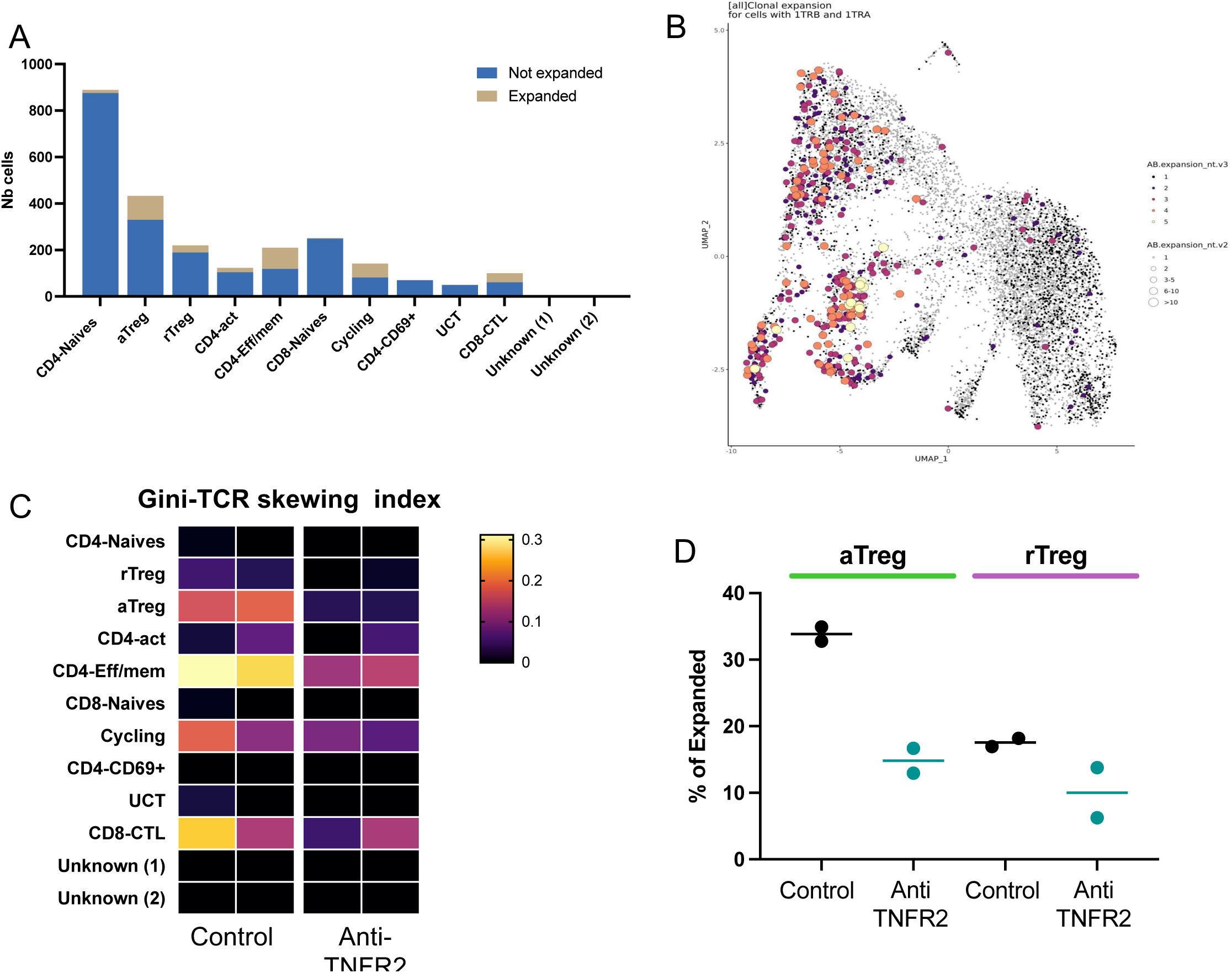
TNFR2 blockade specifically reduces clonally expanded aTreg in a mouse model of PDAC. FvB/n immunocompetent mice were grafted with mPDAC cells into the pancreas. Mice were treated with anti-TNFR2 mAb and tumors were harvest and intratumoral CD4^+^ were isolated for single–cell RNA and TCR sequencing. (A) Distribution of cells with expanded clones among the different identified clusters, the clusters with more expanded cells are Treg, CD4-EFF/MEM and CD8-CTL (B) UMAP representation of expanded clones (same conclusion of A) (C) Gini index (explaining the diversity) of all cluster from all mice (near 1 = expansion of clones, near 0= no expansion). (D) specific effect of treatment on cells with expanded clones inside each Tregs cluster.

We then turned our analysis on the effect of the anti-TNFR2 mAb on CD8^+^ T cells. We focused on the cytotoxic CD8^+^ T cells (CD8-CTL) and the naive CD8^+^ T cells (**Figure 5A**). Anti-TNFR2 treatment induced deregulation of 912 genes in CD8-CTL (**Figure 5B**). A significant decrease in the expression of the markers of exhaustion (*Ctl4*, *Tigit*, *Tim3*, *Entpd1, havcr2*) in the CD8-CTL cluster was individually observed and the exhaustion score taking into account all these markers was lower in anti-TNFR2 mAb treated tumors (**Figure 5C**). Importantly, the proportion of CD8^+^ T cells positive for the exhaustion markers CTLA4, PD-1 and TIGIT was shown to be reduced by flow cytometry analysis in the tumors from mice treated by the anti-TNFR2 compared to those from untreated mice (**Figure 5D**).

**Figure 5.**
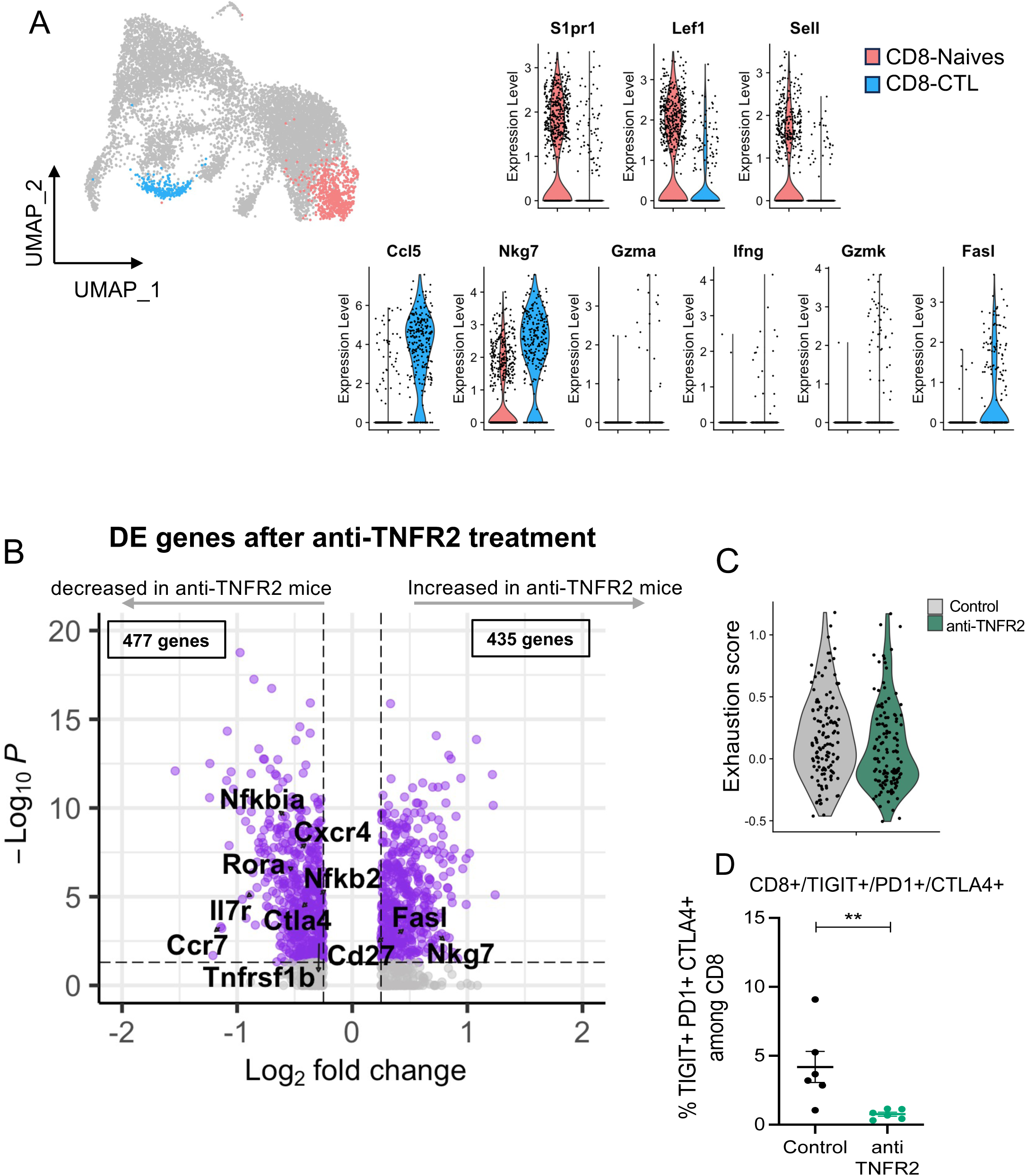
TNFR2 blockade decreases the exhausted profile of CD8 T cells. FvB/n immunocompetent mice were grafted with mPDAC cells into the pancreas. Mice were treated with anti-TNFR2 mAb and tumors were harvest and intratumoral CD8^+^ were isolated for single–cell RNA and TCR sequencing. (A) UMAP plot showing CD8 clusters differentiated by color (CD8-CTL in blue and CD8-Naives in coral) and violin plot comparing expression of different genes between CD8-CTL and CD8-Naives. (B) Volcano plots of differential gene expression analysis between control and anti-TNFR2 treated mice within CD8-CTL cluster. (C) Violin plot showing expression level of CD8 exhaustion score (genes = Tigit, Havcr2, Ctla4, Lag3 and Tox) for each condition (gray: control group, green: anti-TNFR2 treated group). (D) Scatter plot of TIGIT^+^ CTLA-4^+^ PD1^+^ cells proportion among intratumoral CD8^+^ cells analyzed by flow cytometry. Data are plotted as the mean ± SEM. Statistical significance[from controls was determined using a Mann-Whitney test **p< 0,01.

### Combination therapy with blocking anti-TNFR2 and agonistic anti-CD40 mAbs has synergistic effect on PDAC anti-tumor response

We seeked to improve the anti-tumoral response induced by the anti-TNFR2 mAb by designing a combined treatment with immunotherapy that could improve T cell activation. In onco-immunology, agonist mAbs that target costimulatory pathways such as CD40 and OX40 have been shown to succesfully promote antigen-specific T-cell expansion [39,40]. Here, mice were treated with a blocking anti-TNFR2 mAb and agonistic anti-CD40 (**Figure 6A**) administered at day 11, 13, 15. Twenty one days after PDAC cell injection, mice were euthanized and the tumor volumes were measured (**Figure 6B**). Whereas the administration of an anti-TNFR2 mAb or an anti-CD40 agonist alone tends to reduce tumor progression, only the anti-TNFR2 mAb combined with an agonistic anti-CD40 mAb (but not with an agonistic anti-OX40, data not shown) significantly reduced tumor growth (**Figure 6B**). Infiltrated T cell proportion were analyzed by flow cytometry. Whenever agonist anti-CD40 mAb was administered, alone or in the presence of anti-TNFR2 mAb, an increase in CD3^+^, CD8^+^CD4^+^ and Foxp3^-^ T cells (**Figure 6C-F**), and a decrease in Treg (**Figure 6G**) were observed, in line with an improved anti-tumor immune response. Importantly, only the combined treatment induced higher frequency of IFNγ or Granzyme B expressing T cells compared to anti-TNFR2 mAb alone (**Figure 6J-L**). However, in a dedicated experiment, we did not observed improved survival of mice whatever the treatment used compared to untreated mice (**Supplementary Figure 5**).

**Figure 6.**
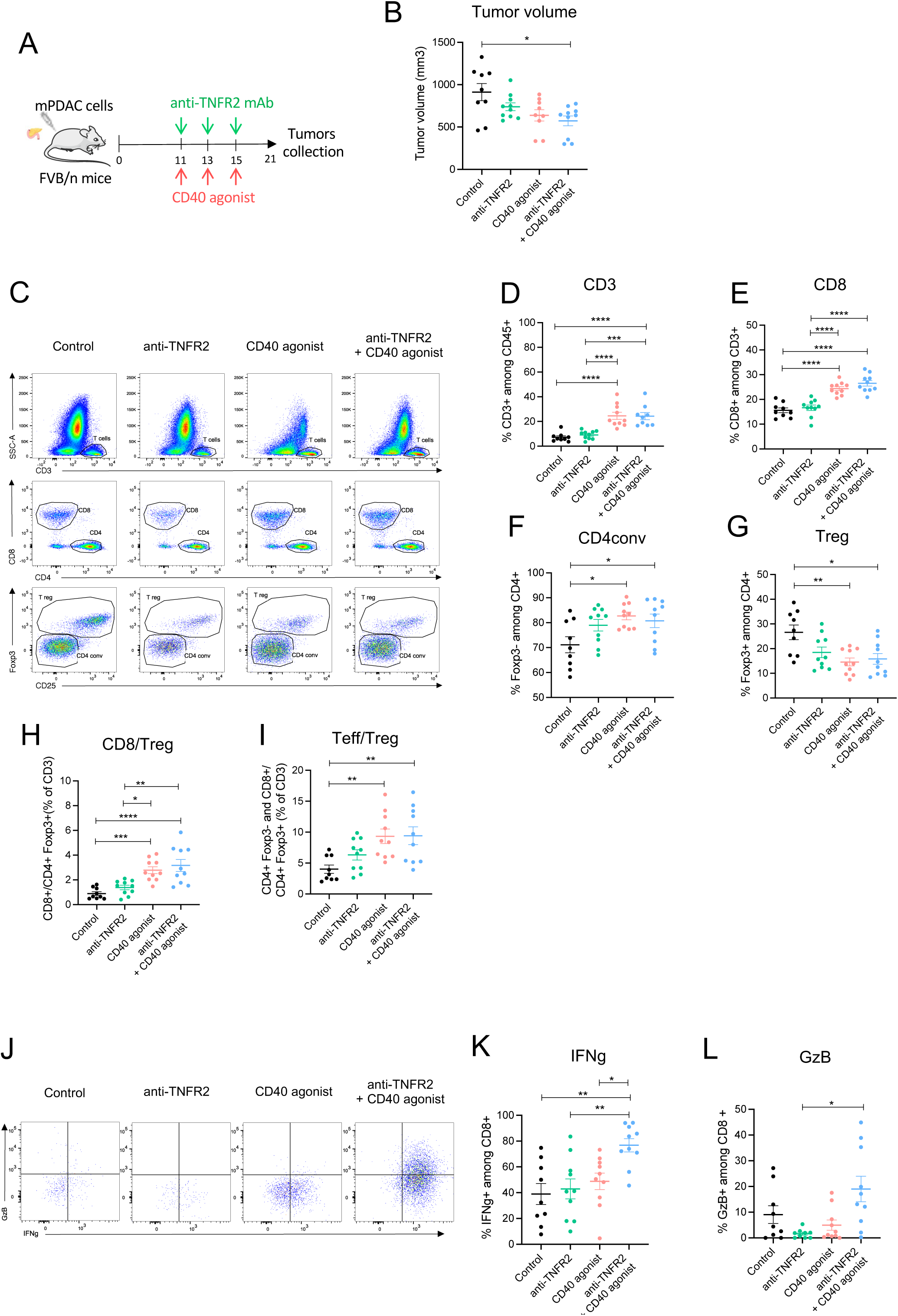
Combination of blocking anti-TNFR2 and agonistic anti-CD40 mAbs increases effector T cell activation. **(A)** Schemas of the experiment: FvB/n immunocompetent mice were grafted with mPDAC cells into the pancreas. Mice were treated with either anti-TNFR2 mAb or CD40 agonist or both or received PBS at day 11, 13, 15 (n=10). After 21 days, tumors were harvest for flow cytometry analysis **(B)** Scatter plot of tumor volume at day 21. **(C-G)** Scatter plots of intratumoral CD3^+^, CD8^+^, Treg (CD4^+^ Foxp3^+^) and CD4conv (CD4^+^Foxp3^-^) cells proportion with representative dot plot (**C**). **(H-I)** Scatter plots of CD8^+^/Treg(CD4^+^Foxp3^+^) ratio and Teff (CD4^+^Foxp3^-^ and CD8^+^)/Treg(CD4^+^ Foxp3^+^) ratio. **(J)** Representative dot plot of IFNg^+^, and GzB^+^ (granzyme B) cells proportion in intratumoral CD8. **(K-L)** Scatter plots of IFNg^+^ and GzB^+^ proportion in CD8 cells. Data are plotted as the mean ± SEM. Statistical significance from controls was determined using a Kruskal-Wallis test [B-N] or ANOVA (depending of the data following a normal distribution), ns: non-significant p>0,05, *p<0,05, **p< 0,01, ***p<0,001, ****<0,0001.

In order to confirm our biological and clinical observations obtained with the combined treatment, we reproduced these experiments using the Panc02-derived orthotopic tumors (**Figure 7A**). The treatments between mPDAC and Panc02 tumors for evaluating tumor volume were similar (**Figure 6A and 7A**). After 21 days, mice were euthanized. As observed for mPDAC, the combination treatment had a statistically significant impact on the tumor incidence compared to control and anti-TNFR2-treated mice (**Figure 7B**). Remarkably, 70% of mice treated by the combination treatment versus 50% treated by anti-CD40 did not develop tumors (**Figure 7B**). The tumor mass of mice treated with the combination treatment that still developed tumors was very low compared to control, anti-TNFR2 or anti-CD40 treated mice (**Figure 7C**). We previously described that mPDAC bearing mice die between the third and the fourth week after cancer cell injection into the pancreas [28]. In a second set of experiments, we assessed whether the observed antitumor effects of anti-TNFR2, anti-CD40 or the coadministration of both mAbs would also enhance the survival of Panc02- bearing mice. Whereas Panc02 control mice died after the fourth week, the anti-CD40 and the anti-TNFR2 mAbs induced a statistically significant improved survival. The better survival was obtained in mice receiving the combined treatment (**Figure 7E**). Strikingly after 64 days, 55% of mice treated with anti-TNFR2 and anti-CD40 mAbs were still alive and tumor free compared to 20% mice with the sole anti-CD40 (**Figure 7F**). In protected mice, we evaluated the possible presence of sub-clinic tumors by ultrasound. Of the 13 mice treated with the Ab combination, tumors were detected in 2 of them. One third of the anti-CD40-protected mice had tumors.Then, we rechallenged the alive mice from the combination group with Panc02 cells injected subcutaneously (**Figure 7G**). Control mice developed subcutaneous tumors as in Supplementary Figure S1 while the anti-TNFR2/anti CD40 mice did not have visible tumors growing suggesting that these mice rejected Panc02 cells due to acquired immunological memory (**Figure 7H**).

**Figure 7.**
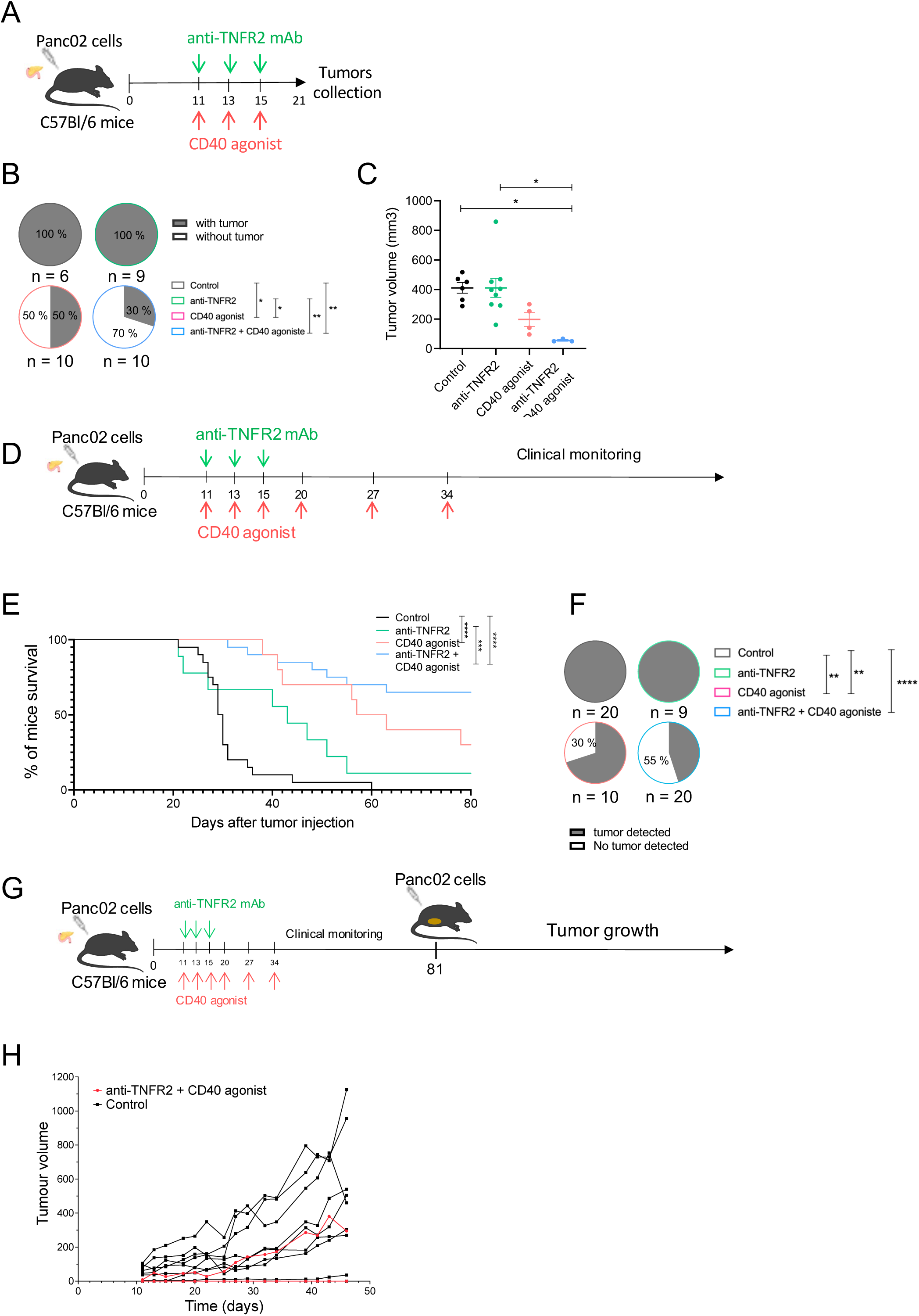
Combination of blocking anti-TNFR2 and agonistic anti-CD40 mAbs induces immunological T memory and improves survival of mice. **(A)** Schemas of the experiment: C56Bl/6j immunocompetent mice were grafted with Panc02 cells into the pancreas. Mice were treated with either anti-TNFR2 mAb or CD40 agonist or both or received PBS at day 11, 13 and 15 (n=10). Mice were euthanized at day 21. **(B)** Proportion of tumor incidence in control and treated groups represented in pie charts (Chi square, * p<0,05, **p<0,01) **(C)** Scatter plot of tumor volume at day 21. Data are plotted as the mean ± SEM. Statistical significance from controls was determined using a Kruskal-Wallis test. *p<0,05. **(D)** Schemas of the experiment: C56Bl/6j immunocompetent mice were grafted with Panc02 cells into the pancreas. Mice were treated with either anti-TNFR2 mAb or CD40 agonist or both or received PBS at day 11, 13 and 15, 20, 27 and 34 and were clinically monitored. **(E)** Kaplan-Meier survival curve of the following groups: control (n = 20), anti-TNFR2 (n = 9), CD40 agonist (n = 10) and anti-TNFR2+CD40 agonist (n = 20). Mice are euthanized when the limit of the clinical score (established by a grid of symptoms) is reached. (Kaplan-Meier test, *** p<0,001, **** p<0,0001). **(F)** Presence of Tumor (primary and/or metastatic) is challenged by echography at day 64. (Chi-square, ** p<0,01, *** p<0,001, **** p<0,0001). **(G)** Schemas of the experiment: mice still alive from anti-TNFR2 + CD40 agonist group and C57Bl/6 naive mice received subcutaneous injection of Panc02 cells at day 81, and monitored for tumor growth. Tumor growth for individual mice is shown in **(H)**.

## Discussion

In this study, we evaluated the relevance of inhibiting the immunosuppressive effect of Treg by blocking the TNF/TNFR2 pathway in PDAC. We first validated TNFR2 as relevant target by showing that, both in humans and in mice, tumor-infiltrating Tregs express TNFR2 more intensively than other TME TNFR2^+^ cell populations. In mice, treatment with anti-TNFR2 reduced the percentage of Treg in the TME of PDACs, thereby reducing tumor growth, paralleled with a reduction in the percentage of CD8^+^ T cells displaying an exhausted phenotype. Moreover, the combination of the anti-TNFR2 mAb with agonistic anti-CD40 mAbs improved survival in PDAC-bearing mice and promoted immunological memory.

This study corresponds to the first accurate description of TNFR2 expression in PDAC patients by single cell analysis.Using published dataset of single cell RNAseq of 24 tumors from PDAC patients compared to 11 non-PDAC pancreas biopsies [30], we showed that the main cell populations expressing *Tnfrsf1b* in PDAC patients are T cells, macrophages and endothelial cells. The TME is also characterized by the presence of immunosuppressive cells such as Tregs and MDSCs that both require TNFR2 for suppressive functions [41]. Importantly, the cell population that express *Tnfrsf1b* at the highest level in human PDAC turned out to be Treg while *Tnfrsf1b* expression in tumor cells remained at low level. In human PDAC, we thus validated that TNFR2 was a relevant target whose blockade could preferentially impact Treg and to a lesser extent tumor cells. We then focused the rest of our study on evaluating the impact of a TNFR2-blocking treatment on tumor growth and modulation of the immune environment in mouse models of PDAC.

We used our previously described murine model of orthotopic PDAC, in which we had initially observed that the immune TME was similar to that of human PDAC, characterized by an immune-escape profile, enriched in Treg and with a paucity of CD8^+^ T cells [29]. Here, deeper analysis revealed that TNFR2 was expressed by T cells, myeloid cells and to a lesser level by tumor cells. Among T cells, the intensity of TNFR2 expression in Tregs was 3.5 fold higher than in CD4^+^FoxP3^-^ T cells. We therefore have a relevant experimental model to test TNFR2-based immunotherapies due to similarities with what is observed in human PDACs.

Some groups demonstrated that Treg depletion elicits effective anti-tumor immunity in mouse PDAC and supports the efficacity of a potential therapeutic strategy targeting Tregs [15,42]. Different strategies of Treg targeting during PDAC progression have been tested: total Treg ablation in *Foxp3*^DTR^ mice was sufficient to evoke effective anti-tumor response in orthotopically implanted KPC cells (**K**ras G12D and **p**53 R172H mutated cells). This was associated with an induction of cytotoxic CD8^+^ T cells activated by CD11c^+^ dendritic cells [15]. In opposite, anti-CTLA4 or anti-CD25 blocking Abs had no clinical effect in mPDAC [5,42] nor in a transgenic KC^iMist1^ mouse model spontaneously developing PDAC tumors [43], suggesting that total Treg ablation was more effective than treatment by anti-CTLA4 or CD25. Since we observed that TNFR2 was highly expressed by Tregs in human and mouse models of PDAC, blocking TNFR2 was tested as an alternative approach for Treg targeting in a clinically compatible approach using anti-TNFR2 mAb treatment. In two immunocompetent mouse models of orthotopically implanted PDAC, we demonstrated that blocking TNFR2 with the TR75-54.7 mAb was able to reduce tumor infiltrating Treg and promote a more favorable balance between CD4conv/Treg and globally between Teff/Treg cells for anti-tumor response. This result was important if we consider that anti-CTL4A or anti-CD25 antibodies did not show a significant change in CD4^+^Foxp3^+^ Treg in the PDAC tumor [42]. Moreover, compared to anti-CTLA4 or anti-CD25 targeting treatments [42], Treg and T cells in LNs did not vary under anti-TNFR2, suggesting that the TNFR2 blockade in our models mainly targets TME-resident Tregs. This could be an important advantage to limit off target effects and potential systemic Treg depletion. These results were reproducible in two different models of orthotopic PDAC and demonstrate that the anti-TNFR2 has a strong and relevant immunomodulatory effect in PDAC. In a recently published study using the same anti-TNFR2 mAb, the depletion of TNFR2 in PDAC murine cells reduced the tumor growth of orthotopic KPC-induced tumors both in immunocompetent or immunodeficient mice, suggesting that TNFR2 expression in tumor cells directly participate to tumor cell growth [44]. Their starting point was the observation of TNFR2 expression in human tumor cells using IHC and single cell analysis. However, at no point was the intensity of TNFR2 expression in tumor cells compared, for example, with that observed in other cell populations of the TME, including Tregs as we did in human PDAC. In our study, in mPDAC and Panc02 orthotopic tumors, TNFR2 blockade impacts tumor growth likely by directly acting on immune cells but not on tumor cells as suggested by the fact that we did not observe any anti-tumor response nor immunomodulatory effect in ectopic compared to orthotopic tumors generated by the same PDAC cells. In our hands and using two different PDAC models, the pancreatic immune TME was sensitive to the treatment and at the center of the clinical effect observed.

We then tried to better define the mechanism of the TNFR2 inhibition by single cell transcriptomic analysis of T cells infiltrated in the TME in untreated and treated mice. We found that Tregs within mPDAC TME were clustered in two groups: aTregs and rTregs. These clusters were identified based on a similar gene expression signature as the one found in physiological condition [45]. Interestingly, the signature found in the aTreg cluster from the pathological murine PDAC TME includes genes hindering tumor immunity, which were previously indentified in Treg from PDAC patients [46]. Indeed, highly immunosuppressive states of Tregs in PDAC patients were associated with high *Tigit*, *Icos* and *Cd39* expression. The anti-TNFR2 mAb treatment strongly impacted the cluster of aTregs compared to rTregs by decreasing specific markers. In particular, a signature of genes associated to the activated Treg clusters *Tnfrsf1b*, *Ctla4*, *Icos*, *Cd39, Il2ra*, *Tigit*, *Tnfrsf4*, *Tnfrsf9*, *Ccr8*, *Pdcd1* significantly decreased in the PDAC TME of mice treated with anti-TNFR2 antibody. Using flow cytometry analysis, the frequency of Treg expressing TNFR2, CTLA4 or TNFR2 and CTLA4 was significantly reduced after anti-TNFR2 treatment, thus validating the transcriptomic data.

CD8^+^ T cells in human PDAC have an exhausted and senescent profile [46] and phenotypes of exhaustion were clearly identified in the TME of our mouse models of PDAC. TNFR2 blockade did not change the phenotype of naïve CD8^+^ T cell cluster, but impacted the expression of exhaustion markers of CD8-Eff/mem T cells. A transciptomic exhaustion score calculated by the expression of *Tigit*, *Lag*3, *Pdcd1*, *Entpd1*, *Ctla4* and *Havcr2,* and the frequency of CD8^+^ T cells expressing TIGIT, PD1 and CTL4A were significantly decreased upon anti-TNFR2 treatment. TNFR2 inhibition is therefore able to affect Treg activation and CD8^+^ T cell exhaustion in the TME. Morevoer, TCR sequencing revaled that TNFR2 blockade decreased the frequency of expanded aTreg clones.

Finally, TNFR2 blockade alone was not sufficient to decisively boost the clinical anti-tumoral response. However, the strong restriction of activated Tregs and the repogramming of naïve T cells suggested that a combination therapy with an activator of CD8^+^ T cells could promote more efficient T cell responses against PDAC. Importantly, we showed that the combined treatment associating anti-TNFR2 and an agonistic anti-CD40 mAbs was able to increase CD3^+^ and CD8^+^ T cell frequency, as well as the expansion of IFNγ- and granzyme B-producing CD8^+^ T cells. This combined treatment impacted tumor growth in mPDAC mouse models compared to control mice and to mono-therapies with the anti-TNFR2 or anti-CD40. Moreover, the survival of mice bearing Panc02-derived tumors was improved compared to control mice and 55% of anti-TNFR2 plus anti-CD40-treated mice were tumor free and developped a memory response against tumor re-challenge in the absence of any additional treatment. Targeting immune checkpoints that suppress anti-tumor immune responses, mainly by using anti-CTLA4 and anti-PD-1/PD-L1 monoclonal antibodies (mAb), has demonstrated robust clinical activity in several cancers, but not in PDAC [5]. Recently, the CD40 agonist antibody tested in a phase II clinical trial did not show benefit [47]. However, despite the failure of this immunotherapy, some patients had increased intratumoral T cell infiltration. Here, we provide a proof of principle in mice that treatment combining a CD40 agonist and a TNFR2 antagonist represents a novel promising immunotherapy approach that deserves to be tested to effectively treat PDAC patients.

## Author’s disclosures

E.P. is co-founder and consultant for Egle-Tx.

## Author contributions

JLC, IC, EP and BLS designed the study; AD, CP, SM, OB AT, CH, MP, JM and NHT performed experiments; AD, CP, PC, JTB ans SB performed mouse single cell experiments, AD, CP, SM, OB, AT, EP, IC, and JLC analyzed the data; CP, WR, EP, analysed single cell data, BLS edited the manuscript, AD, CP, EP, IC and JLC wrote the manuscript.

## ACKNOWLEDGEMENTS

Anaïs Debesset received a PhD grant from the Université Paris-Est-Créteil (UPEC). Sylvain Fisson was supported by grants from the National Institute for Cancer Research (INCA PRT-K 18-022 “IMPROVE ICI”). This work was supported by INCA PRT-K 18-022 “IMPROVE ICI”, ANR-Theranuc from the Agence Nationale de la Recherche and the French charitable organisation “Ligue National contre le Cancer”. We are grateful to the IMRB for providing access to their animal facility team and the flow cytometry platform team for their help. We thank Virginie Reynal, Benoit Albaud, Laura Baudrin, and Patricia Legoix at the Curiecoretech Next Generation Sequencing (ICGEX) platform at Institut Curie. High-throughput sequencing was performed by the ICGex NGS platform of the Institut Curie supported by the grants ANR-10-EQPX-03 (Equipex) and ANR-10-INBS-09-08 (France Génomique Consortium) from the Agence Nationale de la Recherche (“Investissements d’Avenir” program), by the ITMO-Cancer Aviesan (Plan Cancer III) and by the SiRIC-Curie program (SiRIC Grant INCa-DGOS-465 and INCa-DGOSInserm_12554). Data management, quality control and primary analysis were performed by the Bioinformatics platform of the Institut Curie. We acknowledge Coralie Guerin, Anna Chipont, Annick Viguier, and Lea Guyonnetin at Curiecoretech Cytometry plattform at Institut Curie. The authors are grateful to Dr. Sebastien Lemoine for his decisive help with the analysis in humans in sRNA-seq data. This work was supported by the LabEx DCBIOL (ANR-10-IDEX-0001-02 PSL; ANR-11- LABX-0043); and Center of Clinical Investigation (CIC IGR- Curie 1428).

## Supplementary Figures

**Supplementary Figure 1. TNFR2 blockade effects in Panc02 orthotopic mouse model.**

C57Bl/6 immunocompetent mice were grafted with Panc02 cells into the pancreas. After 21 days, tumors were harvest for flow cytometry analysis. **(A)** TNFR2 expression in intratumoral Treg (CD4^+^ Foxp3^+^), CD4conv (CD4^+^ Foxp3^-^) and CD8^+^ cells **(B)** Histograms show TNFR2 expression (MFI TNFR2) and **(C)** proportions among intratumoral Treg (CD4^+^ Foxp3^+^), CD4conv (CD4^+^ Foxp3^-^) and CD8^+^ cells are shown in the graphs (n=6). **(D)** Schema of the experiment: mice were treated with anti-TNFR2 mAb or PBS at days 8,11,13 and 15 (n=6). **(E)** Scatter plot of tumor volume at day 21. **(F-G)** Scatter plots of intratumoral Treg (CD4^+^ Foxp3^+^), and CD8^+^ proportions. (H) TNFR2 proportion among CD4^+^Foxp3^+^. **(I-J)** Scatter plots of CD8^+^/Treg(CD4^+^ Foxp3^+^) ratio, and Teff (CD4^+^Foxp3^-^ and CD8^+^)/Treg(CD4^+^Foxp3^+^) ratio. **(K)** Schema of the experiment: FvB/n immunocompetent mice were grafted with mPDAC cells subcutaneously in the right flank and were treated with anti-TNFR2 mAb or IgG control at days 8, 11 and 13 (n=10). After 20 days, tumors were harvest for flow cytometry analysis. **(L)** Tumor growth curve and **(M)** tumor volume at day 20 are shown in the graphs. **(N)** Scatter plot of intratumoral Treg (CD4^+^ Foxp3^+^) proportion. Data are plotted as the mean±SEM. Statistical significance between population in control was determined using Kruskal-Wallis test, *p<0,05 **p<0,01. Statistical significance from controls was determined using Mann-Whitney test (two-tailed [H-K-N-O] or one-tailed [E]. ns: non-significant p>0,05, *p<0,05, **p< 0,01.

**Supplementary Figure 2. TNFR2 blockade does not modify T cells proportions in tumor draining lymph nodes.**

FvB/n mice injected orthotopically with mPDAC cells were treated with anti-TNFR2 mAb, or IgG control, PBS at day 11, 13 and 15 or were untreated. After 21 days, pancreatic draining lymph nodes were harvest for flow cytometry analysis. **(A-B)** Scatter plots of Treg (CD4^+^ Foxp3^+^) and CD8^+^ cells proportions **(C-D)** Scatter plots of CD8^+^/Treg(CD4^+^Foxp3^+^) ratio and Teff(CD4^-^Foxp3^+^ and CD8^+^)/Treg(CD4^+^Foxp3^+^) ratio (n=26 including 2 experiments, representatives of 3 experiments). Data are plotted as the mean±SEM. Statistical significance from controls was determined using Mann-Whitney test. ns: non-significant.

**Supplementary Figure 3. TNFR2 blockade effects on myeloid cells.**

FvB/n immunocompetent mice were grafted with mPDAC cells into the pancreas. After 21 days, tumors were harvest for flow cytometry analysis. **(A)** Gating strategy used for myeloid cells identification. **(B)** Histograms of TNFR2 expression and **(C)** Representative histograms of TNFR2 expression (MFI TNFR2) and **(D)** proportions in intratumoral M-MDSC (CD11b^+^Ly6C^+^Ly6G^-^), PMN-MDSC (CD11b^+^ Ly6C^low^Ly6G^+^), Macrophages (CD11b^+^Ly6G^-^ F4/80^+^), M1 (F4/80^+^CD11c^+^CD206^-^), M2 CD206^+^ (F4/80^+^ CD11c^−^ CD206^+^), M2 CD206^-^(F4/80^+^ CD11c^−^CD206^-^) DC: dendritic cells (CD11c^hi^ IA-IE^+^ F4/80^-^) and DC type 1 (CD11c^hi^ IA-IE^+^ F4/80^-^ CD11b^-^ CD103^+^) and DC type 2 (CD11c^hi^ IA-IE^+^ F4/80^-^ CD11b^+^ CD103^-^) (n=6). Mice were treated with anti-TNFR2 mAb, IgG control at day 11, 13 and 15 or were untreated (n=21). After 21 days, tumors were harvest for flow cytometry analysis. **(E-M)** Scatter plots of cell proportion among CD45+ FVS- cells: (E) CD11b^+^, (F) M-MDSC (CD11b^+^ Ly6C^+^ Ly6G^-^), (G) PMN-MDSC (CD11b^+^ Ly6C^low^ Ly6G^+^), (H) macrophages (CD11b^+^ Ly6G^-^ F4/80^+^), (I) M1 (F4/80^+^ CD11c^+^ CD206^-^), (J) M2 (F4/80^+^ CD11c^−^ CD206^+^), (K) DC (CD11c^hi^ IA-IE^+^ F4/80^-^), (L) DC1 (CD11c^hi^ IA-IE^+^ F4/80^-^ CD103^+^ CD11b^-^) and (M) DC2 (CD11c^hi^ IA-IE^+^ F4/80^-^ CD103^-^ CD11b^+^) **(N)** Representative histograms of CD80 (n=21) and CD86 (n=11) expression (MFI CD80 and MFI CD86) on CD11c^+^ IA-IE^+^ (DC like) cells and respective scatter plots **(O-P).** Data are plotted as the mean±SEM. Statistical significance between population in control was determined using Kruskal-Wallis test, **p<0,01 ***p<0,001. Statistical significance from controls was determined using t-test [E-H-J-K-L-M] or Mann-Whitney two-tailed [F-G-J-O-P]. ns: non-significant p>0,05, *p<0,05, **p< 0,01, ***p<0,001.

**Supplementary Figure 4. Identification of the cell clusters from single cell RNAseq analysis of PDAC.**

Intratumoral CD4^+^ and CD8^+^ were isolated for single–cell RNA sequencing from mPDAC control and anti-TNFR2 treated mice. (A) Heatmap showing mean gene expression of different genes used to describe CD4 and CD8 T cells cluster (clustering done using FindNeighbors on the fifty first PCs and FindClusters, resolution 0.7).

**Supplementary Figure 5. Combination of blocking anti-TNFR2 and agonistic anti-CD40 mAbs effects in survival of mPDAC orthotopic mouse model.**

FvB/n immunocompetent mice were grafted with mPDAC cells into the pancreas. Mice were treated with either anti-TNFR2 mAb or CD40 agonist or both or received PBS at day 11, 13, 15, 20 and 27. Clinical score of the mice is established by a grid of symptoms. Mice are euthanized when the limit of the clinical score is reached. **(A)** Kaplan-Meier survival curve of the following groups: Control (n = 20), anti-TNFR2 (n = 10), CD40 agonist (n = 10) and anti-TNFR2+CD40 agonist (n = 20). (Kaplan-Meier test, ns p>0,05).

## Supplementary Materiel

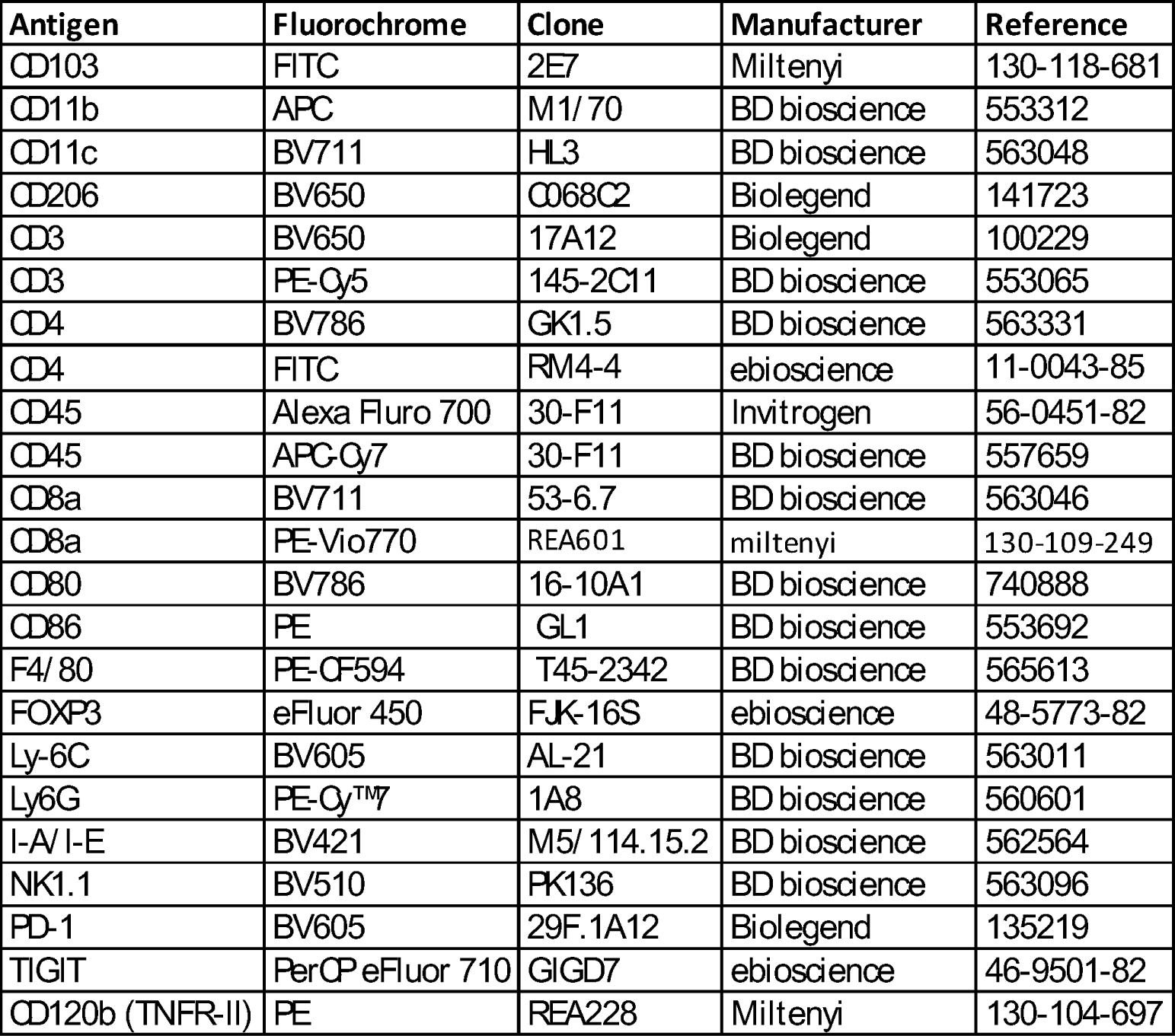
EXTRACELLULAR.

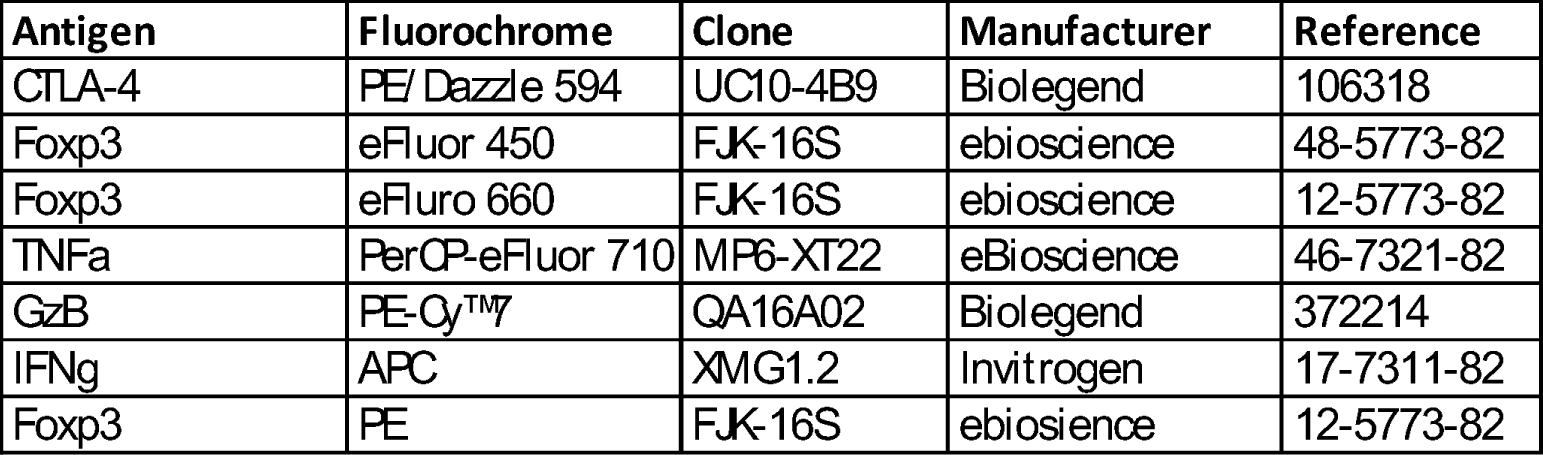
INTRACELLULAR.

FIXABLE VIABILITY STAIN (FVS)

**FIXABLE VIABILITY STAIN (FVS)**
| Antigen | Fluorochrome | Clone | Manufacturer | Reference |
| --- | --- | --- | --- | --- |
| FVS | BV510 |  | BD bioscience | 564406 |
| FVS | Alexa F700 |  | BD bioscience | 564997 |

