## Supplemental material for "TNFR2 blockade promotes anti-tumoral immune response in PDAC by targeting activated Treg and reducing T cell exhaustion"

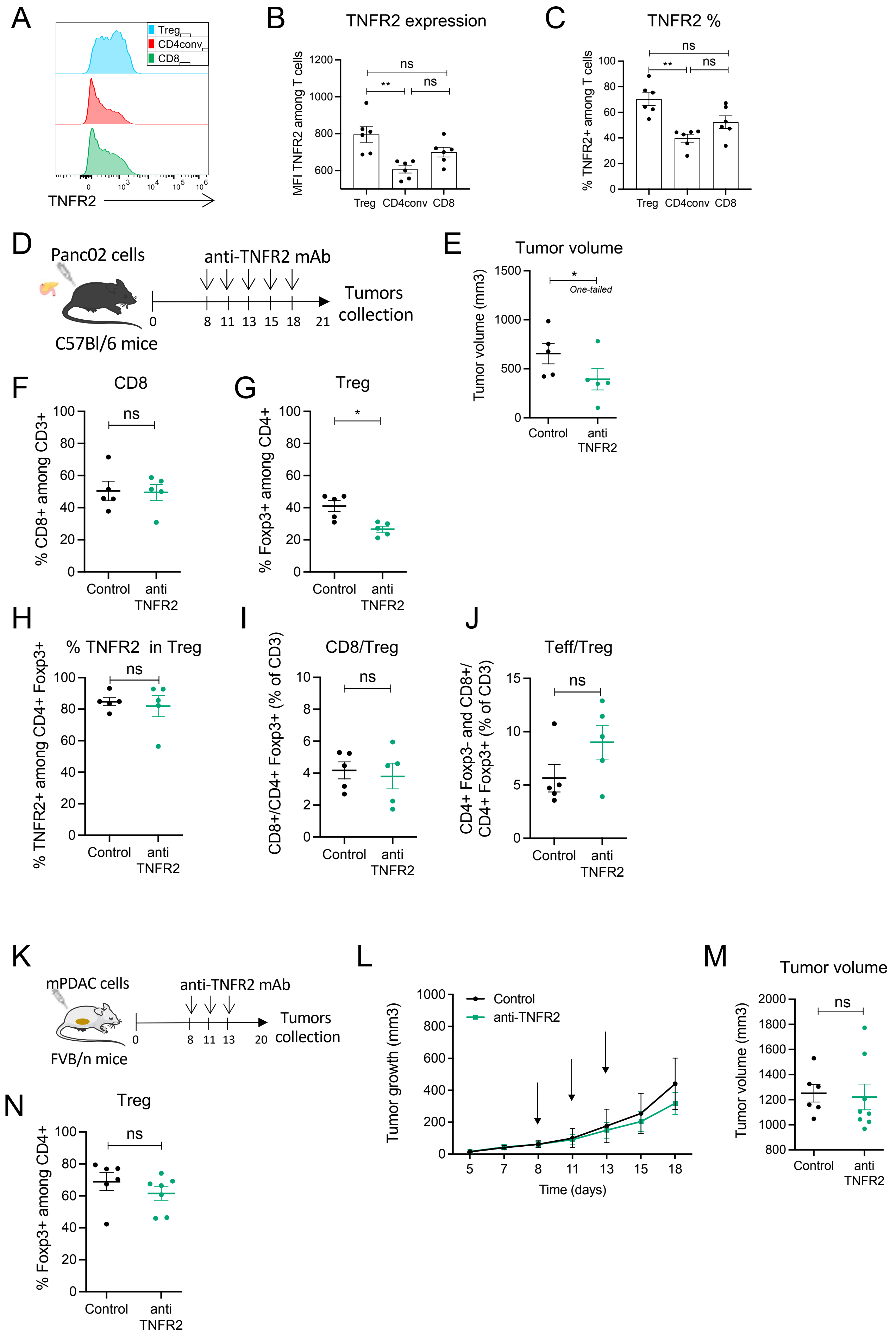


**Supplementary Figure 1**

### Draining lymph nodes

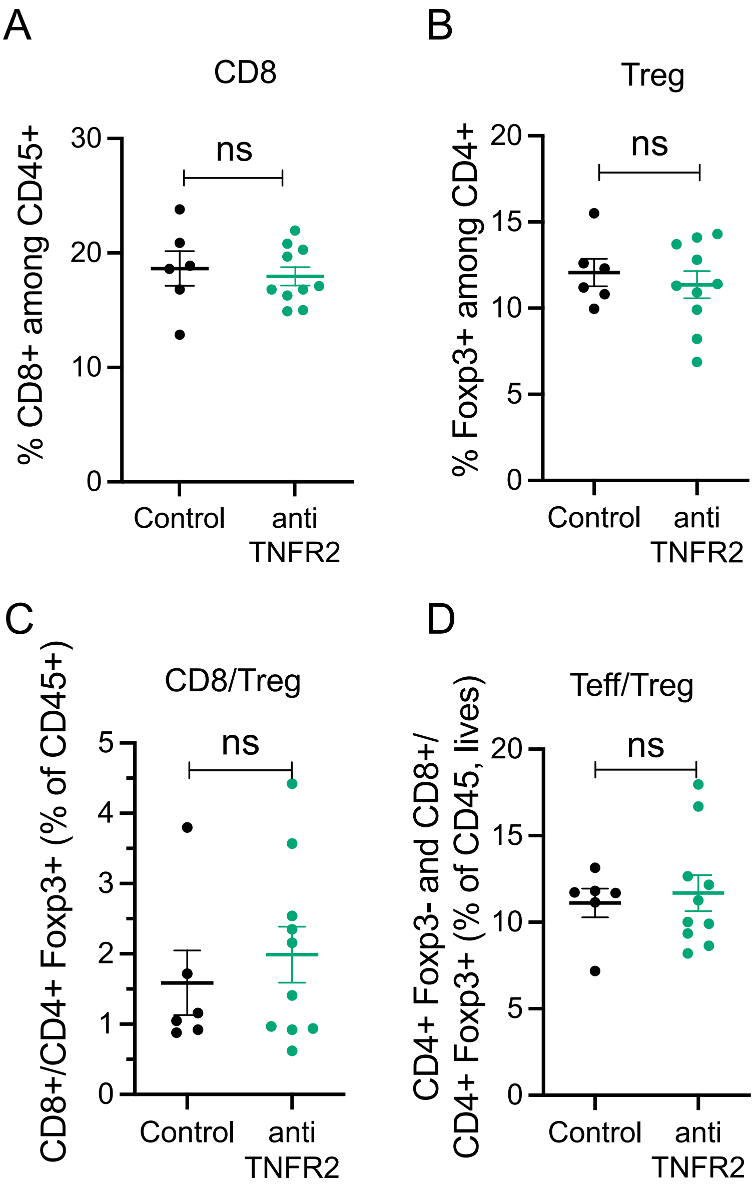


**Supplementary Figure 2**

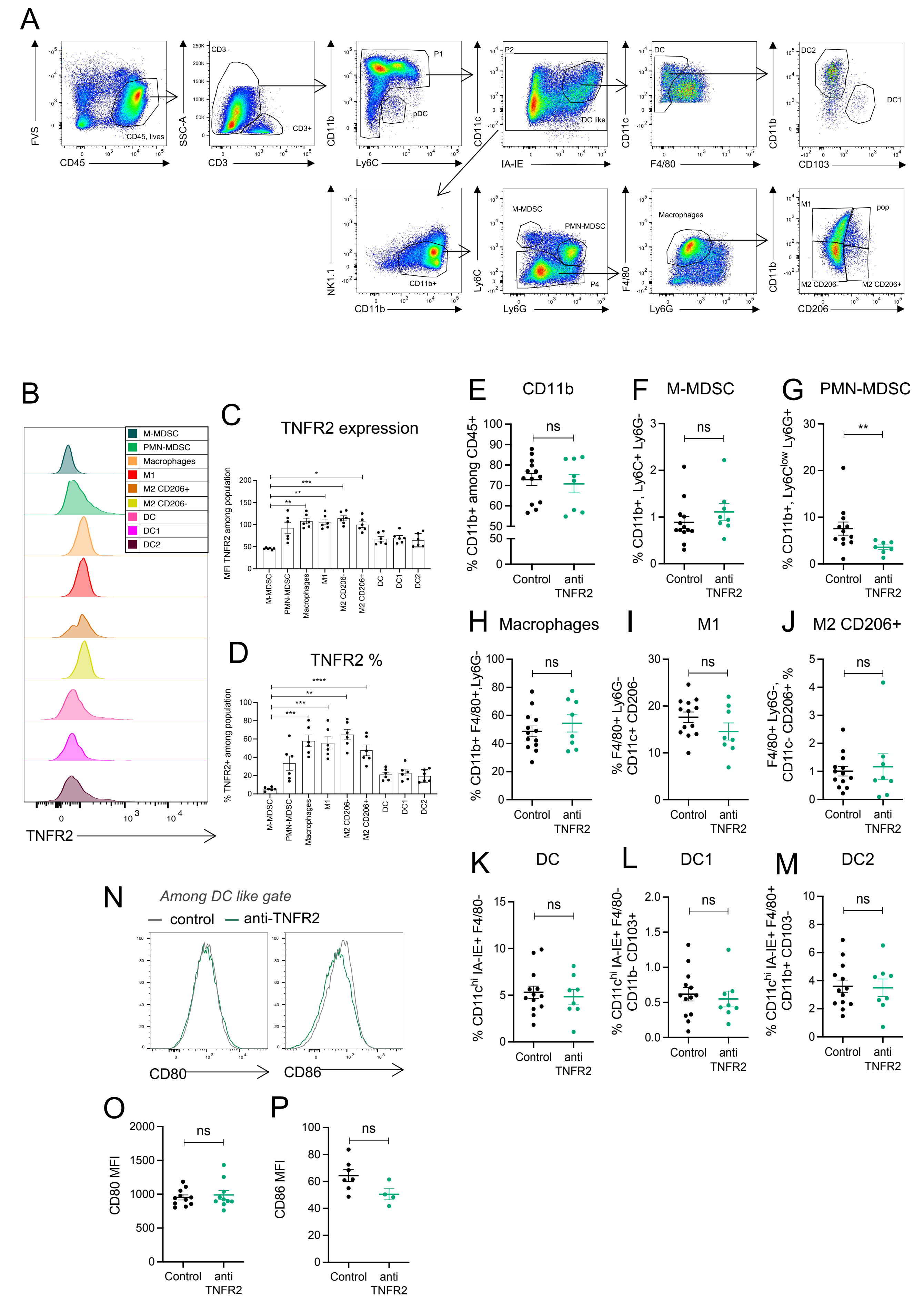


**Supplementary Figure 3**

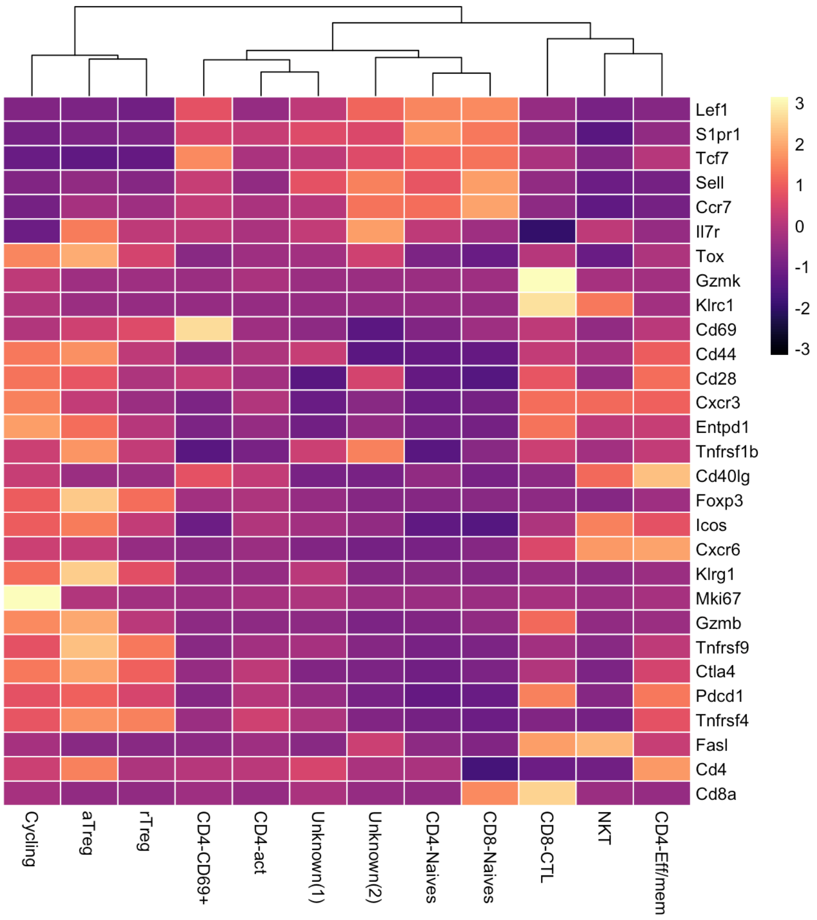


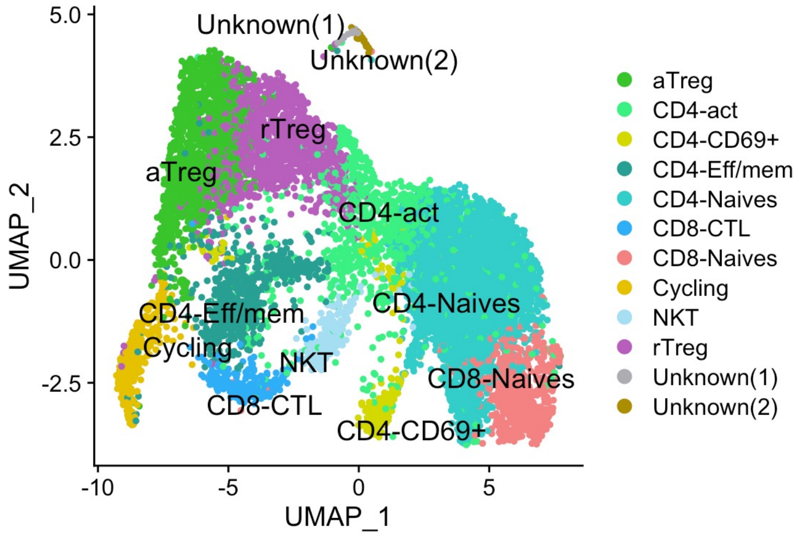


**Supplementary Figure 4**

A

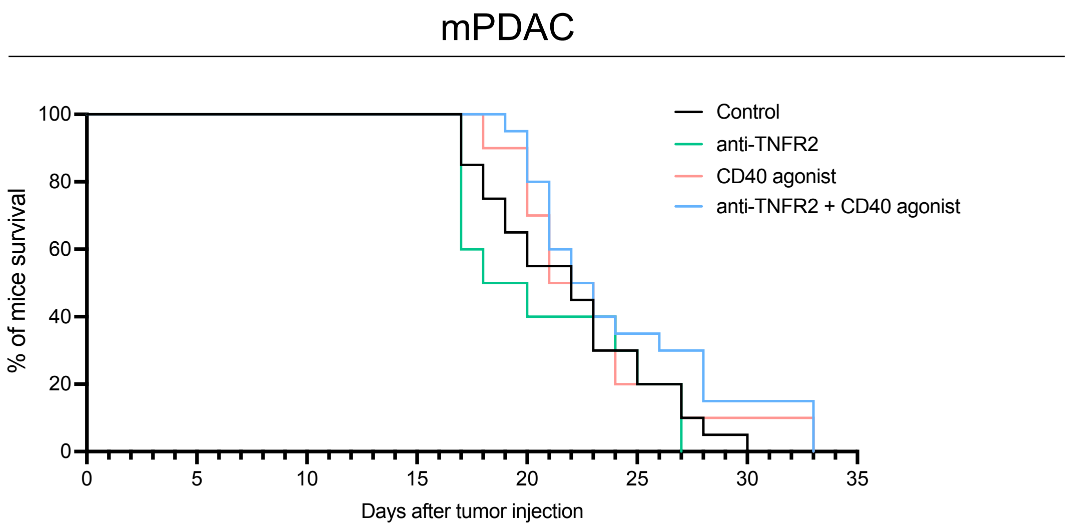


**Supplementary Figure 5**
